# Molecular mechanisms regulating PDE11A4 age-related liquid-liquid phase separation and its reversal by selective PDE11A4-targeted small molecules in cells and brain

**DOI:** 10.1101/2025.05.20.654583

**Authors:** Elvis Amurrio, Janvi Patel, Marie Danaher, Helen Do, Laila Elzaree, Janet L. Fisher, Haley Greene, Porschderek Kargbo, Priscilla Kim, Eliska Klimentova, Sonia Lin, Yingchen Liu, M. Yusuf Mehboob, Suad Abdella, Yadiel Temesgen, Wenbo Yu, Shams Ul Mahmood, David P. Rotella, Michy P. Kelly

## Abstract

Age-related increases in hippocampal PDE11A expression occur in humans and rodents, causing age-related cognitive decline of social memories. Interestingly, this age-related increase triggers PDE11A4 liquid-liquid phase separation (LLPS), causing the enzyme to condensate in neurons forming filamentous structures termed “ghost axons”. Here we sought to identify molecular mechanisms regulating PDE11A4 LLPS and its reversal by therapeutic agents. PDE11A4 LLPS was increased by broad kinase inhibition and overactivation of the trans-Golgi network (via TGN38 or RhoB) but suppressed by PDE11A4-pS163, PDE11A4-pS239 or a D355A mutation affecting allosteric cGMP-GAF-A interactions. PDE11Ai inhibitors across three scaffolds rapidly reversed PDE11A LLPS independently of their catalytic effects, pS163 or pS239, but rather—at least in part—by reducing PDE11 homodimerization at the GAF-B domain. PDE11Ai’s blocked exacerbated LLPS caused by age-related PDE11A4 phosphorylation, kinase disruption, and altered proteostasis in hippocampal HT22 neuronal cells. Further, in hippocampus and amygdala of old mice, oral SMQ-03-20 reversed age-related increases in PDE11A4 condensation in ghost axons and protein expression of immunoglobulin light chains (a marker of increased blood-brain permeability), with direct correlation between the two measures. These findings suggest PDE11A4 LLPS is a regulated and druggable mechanism contributing to age-related hippocampal pathology.\

## INTRODUCTION

Aging is accompanied by progressive cognitive decline, for which there are no approved pharmacological interventions. Liquid–liquid phase separation (LLPS)—the formation of dynamic, membraneless biomolecular condensates—is a biophysical mechanism increasingly recognized as central to proteostasis and is emerging as a key player in the pathology of aging and neurodegeneration ^4–11^. Under homeostatic conditions, LLPS governs the dynamic assembly of biomolecular condensates that regulate protein sequestration, buffering, and clustering with substrates ^8^. However, dysregulated LLPS can seed the formation of aberrant protein aggregates that are now recognized as pathophysiological features of the aging brain ^4–9,11^. This paradigm shift has galvanized intense interest in pharmacological tools that can modulate LLPS (e.g., ^12–15^). Historically, LLPS was considered “undruggable” due to the nature of its inherent biophysical/physiochemical properties, including the involvement of intrinsically disordered regions (IDRs) as well as transient, weak and dynamic interactions ^6,14^. More recently, however, studies have demonstrated that rationally designed or screened small molecules can bind directly to specific intrinsically disordered proteins to alter their phase separation properties and produce a therapeutic benefit (e.g., tau or β-catenin; ^16,17^). Despite this progress, gaps remain regarding generalizability of this approach across different phase separating proteins ^18^.

Phosphodiesterase 11A (PDE11A4) is an enzyme that contains an intrinsically disordered region and undergoes LLPS ^10^. PDE11A is a dual-specificity PDE that hydrolyzes cAMP/cGMP, with the PDE11A4 isoform being the only isoform expressed in neural tissue ^19–22^. PDE11A4 is enriched in the hippocampus, a brain region central to learning and memory ^20,22^. PDE11A4 expression increases with age in both human and rodent hippocampus causing specific hippocampus-dependent memory deficits in mice ^23–25^. Based on these findings, we recently developed the first potent, selective and orally-bioavailable small molecule PDE11A4 inhibitors (PDE11Ai’s) for the treatment of age-related hippocampal pathologies ^1,2,26^.

Interestingly, these age-related increases in PDE11A4 protein expression do not occur uniformly throughout the hippocampus, but rather condensate in filamentous appearing structures named “ghost axons”—that is, axonal terminals that are so densely packed with PDE11A4 protein that other axonal markers are occluded ^24^. Although PDE11A4 ghost axons appear to be continuous filamentous structures at low magnification, higher magnification reveals they are often linear trails of spherical/ovoid puncta ^10,23,24^. The punctate nature of PDE11A4 in the aged brain likely reflects LLPS, as PDE11A4 exhibits multiple sequence features highly predictive of LLPS ^10^ and modeling these aging-like increases in PDE11A4 in HT22 mouse hippocampal cells produces a concentration-dependent emergence of membraneless spherical droplets that progressively fuse over time and are reversed by 1,6-hexanediol, an aliphatic alcohol that interferes with multivalent interactions central to LLPS ^10^. Of greatest relevance to the present study, PDE11A4-targetting molecules that reverse PDE11A4 LLPS *in vitro* (i.e., the small molecule inhibitor tadalafil and the isolated PDE11A4-GAF-B domain) also reverse PDE11A4 ghost axons in the hippocampus of old mice. This supports the idea that PDE11A4 condensation in ghost axons reflects a phase transition rather than aggregation ^10,23,27^. Interestingly, neither cAMP nor cGMP analogs affect PDE11A4 LLPS and cAMP/cGMP-dependent kinases do not block the ability of PDE11A4 inhibitors (PDE11Ai’s) to reverse LLPS, suggesting that the effects of these PDE11Ai’s on PDE11A4 LLPS occur independently of any catalytic effect ^10^.

Although these initial findings were highly intriguing, several gaps in knowledge remained. First, it was not possible to prove that tadalafil reduced PDE11A4 ghost axons by binding to PDE11A4 directly, as opposed to eliciting such effects indirectly via inhibiting PDE5. Traditionally, a compound thought to elicit an effect via PDE11A4 would be tested in *Pde11a* KO mice to ensure the effect of the compound disappeared. In this case, however, the PDE11A4 protein itself was the relevant endpoint. Second, the mechanism by which PDE11Ai’s reversed PDE11A4 LLPS remained unknown. Third, the effect of PDE11A4 ghost axon reversal on a disease-relevant physiological endpoint was not assessed. Here we address these gaps and more by showing (i) the first potent and highly selective PDE11A4 inhibitors reverse PDE11A4 LLPS in vitro—even under conditions mimicking aging or proteostatic stress—and in vivo; (ii) PDE11Ai’s disrupt PDE11A4 LLPS, at least in part, by reducing PDE11A4 homodimerization; and (iii) reversal of age-related PDE11A4 ghost axons by oral administration of the PDE11Ai SMQ-03-20 reduces age-related increases in immunoglobulin light chain accumulation. In addition, we provide a much more in-depth mechanistic understanding of factors regulating PDE11A4 LLPS, thereby identifying novel inroads to therapeutically target PDE11A4 LLPS independent of catalytic inhibitors. These findings suggest PDE11A4 LLPS is a regulated and druggable mechanism contributing to age-related hippocampal pathology.

## RESULTS

Given the hippocampal enrichment of PDE11A4, along with the fact that in vivo effects of compounds will be tested in mice, we employed mouse HT22 hippocampal neuronal cells to model PDE11A4 LLPS as it occurs in aging. In this context, then, in vitro overexpression is not intended to artificially induce condensation, but rather to model the pathological increase in PDE11A4 levels observed in aging. Indeed, this model has proven predictive of in vivo biology, with genetic, post-translational, and pharmacological manipulations that alter PDE11A4 compartmentalization and condensation in cells similarly regulating its compartmentalization and condensation in the aging brain ^10,23,24,27^. This in vitro system thus provides a platform for screening compounds that reverse LLPS and enables the dissection of molecular mechanisms regulating PDE11A4 condensation ^10,24,26^. To facilitate microscopic detection of PDE11A4, we fused emerald-green fluorescent protein (GFP) to the N-terminus of mouse or human PDE11A4. Prior studies confirmed that the addition of the GFP tag does not alter PDE11A4 LLPS, catalytic activity, nor subcellular localization ^10,21,23,24,28^ (for biochemical fractionation, see Figure S1).

### PDE11A4 inhibitors across three chemical scaffolds reverse aging-like PDE11A4 LLPS

To determine whether highly potent PDE11A4-selective small molecules can reverse PDE11A4 LLPS, we tested six of our recently developed “MLG” and “SMQ” series of PDE11Ai’s (Figure 1A) ^1,2^. To enable a direct comparison of potency and efficacy of these six molecules with the two commercially-available compounds we profiled previously ^10^, we also tested tadalafil—a PDE5 inhibitor with potent PDE11A4 inhibiting off-target activity (Figure 1B)—and BC11-38 a low potency PDE11A4-preferring inhibitor (Figure 1C). Notably, these 8 small molecule inhibitors span three chemical scaffolds (Figure 1A-C).

**Figure 1.**
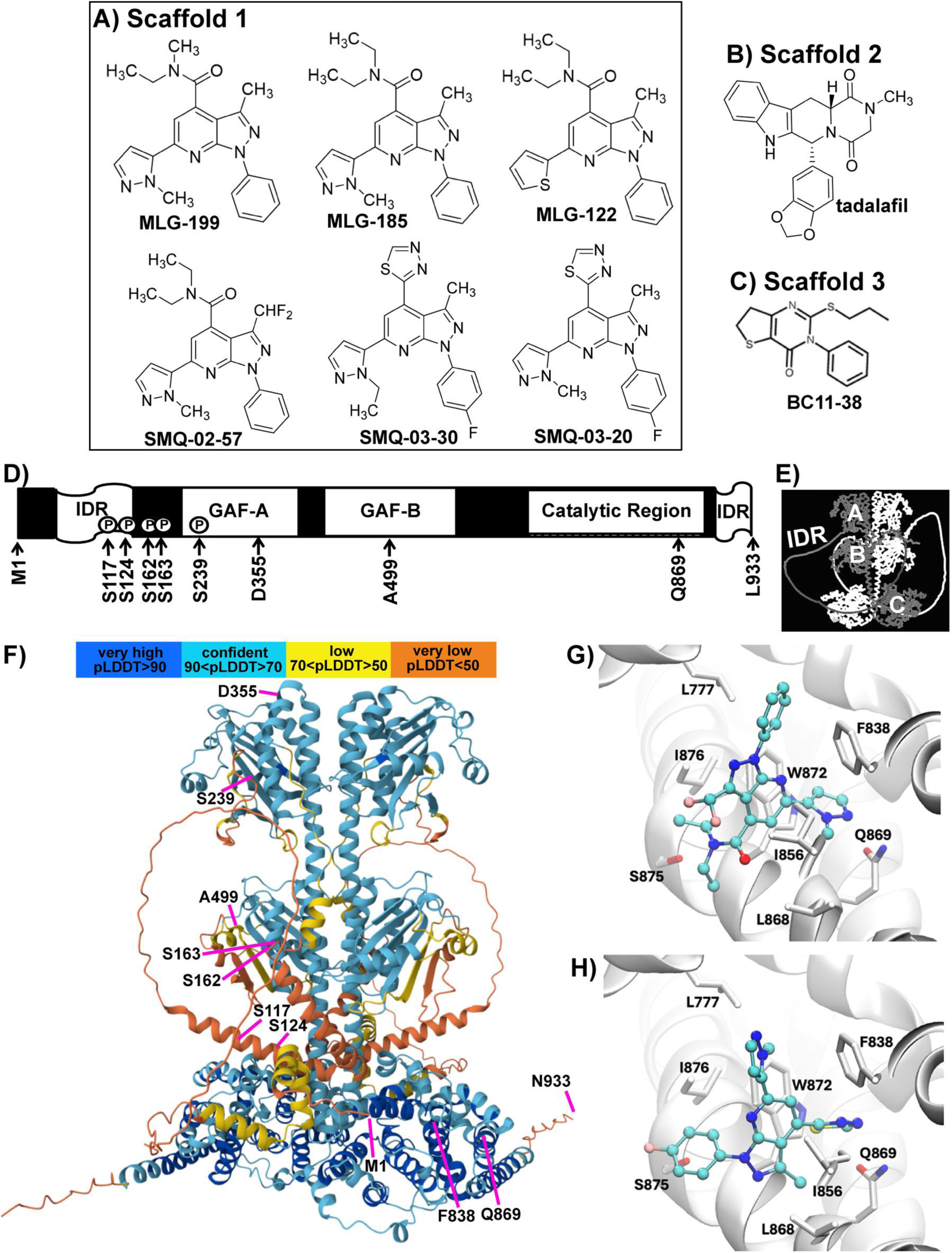
PDE11A inhibitors are predicted to bind the PDE11A4 catalytic domain. Chemical structures for PDE11A inhibitors from A) scaffold 1 (MLG-199, “14k” in 1; MLG-185, “14d” in 1); MLG-122 “9a” in 1; SMQ-02-57, “23b” in 1; SMQ-03-30, “4h” in 2; SMQ-03-20, “4g” in 2), B) scaffold 2 (tadalafil, “1” in 1), and C) scaffold 3 (BC11-38, 3). D) Linearized schematic of a PDE11A4 monomer with notation of key residues, intrinsically disordered regions (IDRs), as well as regulatory (GAF-A, GAF-B) and catalytic domains. E) AlphaFold 2 model of homodimerized PDE11A4 color coded by chain (A—GAF-A domain, B—GAF-B domain, C—catalytic domain). F) AlphaFold 3 model of homodimerized PDE11A4 color coded according to strength of the predicted Local Distance Difference Test (pLDDT) and showing the location of key residues along one of the monomers (see supplement for rotating homodimer with residues noted on both chains). Docking poses of G) SMQ-02-57 and H) SMQ-03-20 shown in Corey–Pauling–Koltun representation. PDE11A4 structure is shown in cartoon representation with binding site residues shown in licorice representations and labeled.

The crystal structure of PDE11A has yet to be solved; however, biochemical and modeling data indicate that tadalafil binds its catalytic pocket with a single hydrogen bond to Q869 being critical for high-affinity binding ^29–32^ (Figure 1D). To determine where our scaffold was likely to bind PDE11A4, we conducted modeling analyses using SMQ-02-57 since it is our most potent PDE11Ai and SMQ-03-20 since it is our only orally bioavailable brain penetrant PDE11Ai to date. First, we used AlphaFold to model the structure of homodimerized PDE11A4 (accessed 05/06/27; Figure 1B-C; see supplemental materials for video of a rotating homodimer). Next, Site Identification by Ligand Competitive Saturation (SILCS) Hotspots analysis was conducted and indicated that the most favorable binding poses for docked SMQ-02-57 (Figure 1G) and SMQ-03-20 (Figure 1H) are located within the well-characterized substrate binding pocket of the catalytic domain. In both cases, the ligands are sandwiched between residues W872 and I856/F838 in the PDE11A4 protein. Notably, the conserved glutamine residue Q869 described above appears to play an important role, with the pyrazolyl group in SMQ-02-57 and the thiadiazole in SMQ-03-20 predicted to form hydrogen-bonding interactions with this residue. The predicted binding affinities of SMQ-03-20 and SMQ-02-57, calculated as ligand grid free energies (LGFE), are −16.49 and −16.75 kcal/mol, respectively, and are consistent with the rank order observed in the experimental IC₅₀ data with SMQ-02-57 being slightly favored (Table S1).

PDE11Ai’s across all three scaffolds rapidly dispersed PDE11A4 LLPS. MLG-199 (Figure 2A-B), MLG185 (Figure 2C), and MLG-122 (Figure 2D) all significantly reduced PDE11A4 LLPS, albeit less potently than did SMQ-02-57 (Figure 2E-F), SMQ-03-30 (Figure 2G) and SMQ-02-30 (Figure 2H). Tadalafil (Figure 2I-J) and BC11-38 (Figure 2K) reduced PDE11A4 LLPS with potencies on par with our MLG series of compounds. These differential potencies across compounds in the LLPS assay correlated with those achieved in the catalytic activity assay (Table S1). PDE11Ai’s also reversed LLPS of GFP-tagged human PDE11A4 in human SH-SY5Y neuronal cells (Figure S2) and prevented mouse PDE11A4 LLPS from occurring in HT22 cells (Figure S3). As previously reported ^10^, and included here as negative controls, PDE11A4 LLPS was not reversed by the PDE10A inhibitor papaverine (Figure 2L) nor the PDE4 inhibitor rolipram (Figure 2M), arguing for specificity of PDE11A4-targeting small molecules as opposed to molecules that increase cAMP and/or cGMP levels. Indeed, we showed previously ^10^ and replicate here (Table S2), that cAMP and cGMP analogues have no effect on PDE11A4 LLPS. Further, we showed previously ^10^ and replicate here (Table S3) that neither cAMP nor cGMP blockers are able to stop the effect of PDE11Ai’s on PDE11A4 LLPS. Further, the effect of our PDE11Ai’s on LLPS are not due to reducing PDE11A4 protein concentrations (Table S1).

**Figure 2.**
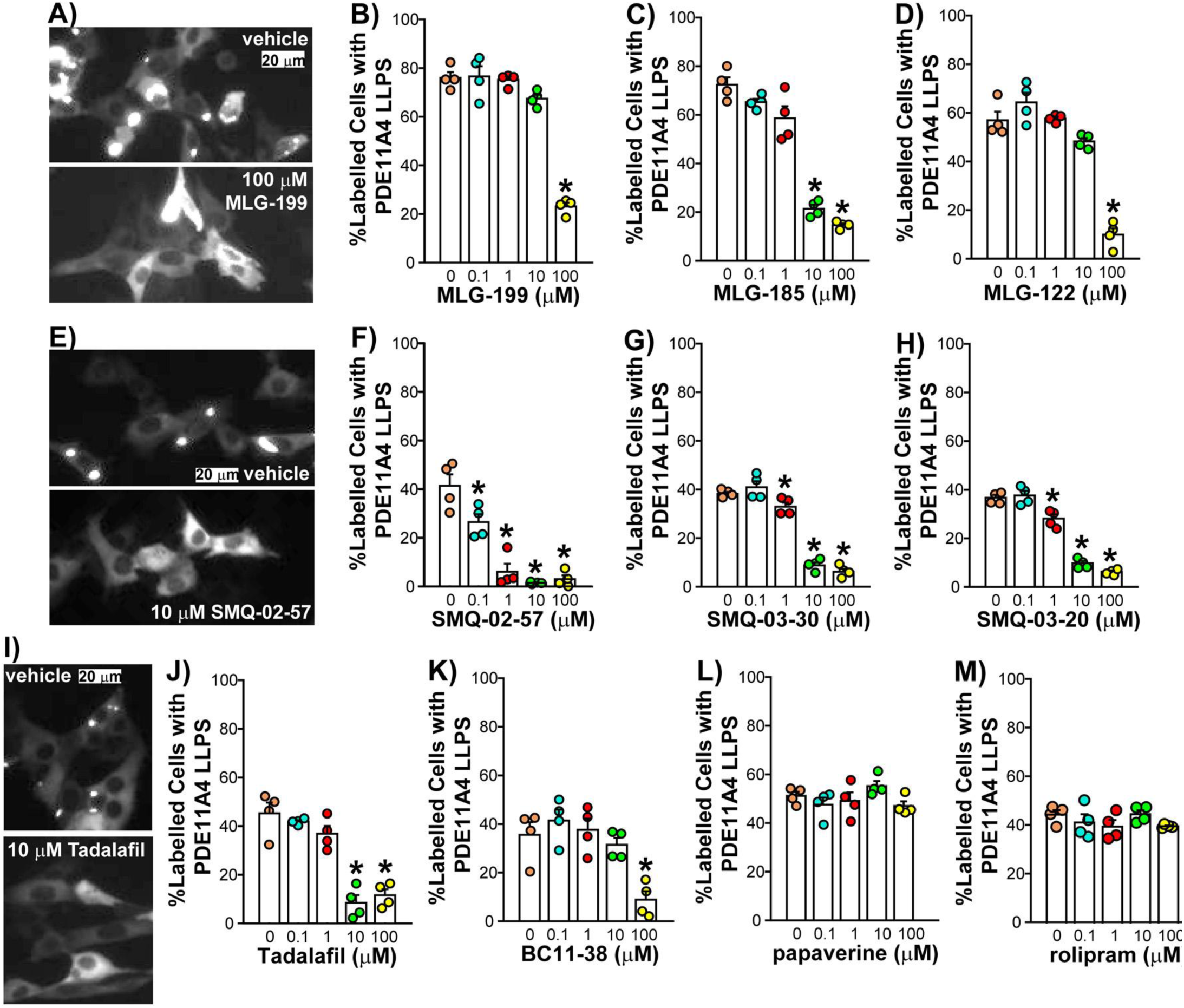
PDE11Ai’s across 3 scaffolds reverse aging-like PDE11A4 LLPS in hippocampal HT22 cells. Mimicking age-related overexpression of PDE11A4 in hippocampal HT22 neuronal cells leads to PDE11A4 LLPS that is reversed by the PDE11Ai’s **A-B)** MLG-199 (ANOVA: F(4,15)=87.43, P<0.0001; Post hoc: 0 µM vs. 100 µM, P=0.0002, **C)** MLG-185 (failed equal variance; ANOVA on Ranks: H(4)=16.4, P<0.0001; Post hoc vs. 0 µM: 1 µM P=0.0102, 10-100 µM P=0.0002 each), **D)** MLG-122 (F(4,15)=61.38, P<0.0001; Post hoc vs. 0 µM: 10 µM P=0.0434, 100 µM P=0.0002), **E-F)** SMQ-02-57 (equal variance fails; ANOVA on Ranks: H(4)=15.61, P=0.0036; Post hoc: 0 µM vs. 0.1-100 µM, P<0.05 for each, **G)** SMQ-03-30 (equal variance fails; ANOVA on Ranks: H(4)=16.74, P=0.0022; Post hoc: 0 vs 1-100 µM, P<0.05 for each), **H)** SMQ-03-20 (fails equal variance; ANOVA on Ranks: H(4)=17.43, P=0.0016; Post hoc: 0 vs 1-100 µM, P<0.05 for each, **I-J)** tadalafil (ANOVA: F(4,15)=32.44, P<0.0001; Post hoc: 0 vs 10-100 µM, P=0.0002 for each), and **K)** BC11-38 (ANOVA: F(4,15)=9.35, P=0.0005; Post hoc: 0 vs 100 µM, P=0.001). In contrast **L)** the PDE10A inhibitor papaverine and **M)** the PDE4i rolipram had no effect on PDE11A4 LLPS. n=4 biological replicates/group. Data are representative of 2 or more experiments *vs. 0 µM, p<0.05-0.0001. Data plotted mean ±SEM.

### Effects of PDE11A4 inhibitors on age-related PDE11A4 LLPS occur within minutes and can reverse rapidly upon washout

Next, we sought to better characterize the pharmacodynamics of the MLG and SMQ PDE11A4 inhibitors. To do so, we treated PDE11A4-transfected HT22 cells for either 1 hour or 24 hours to determine whether prolonged exposure would lead to desensitization, a phenomenon often observed with chronic drug treatment due to compensatory mechanisms. We then washed out the compound for 5 hours to assess whether the inhibitors bind reversibly or irreversibly (Figure S4A). Only MLG-199 (Figure S4C-F), MLG-185 (Figure S4G-J), SMQ-02-57 (Figure S4B and K-N) and SMQ-03-30 (Figure S4O-R) were tested in the washout assay due to limited availability of the other two compounds. Consistent with our previous characterization of tadalafil and BC11-38 ^10^, the MLG and SMQ PDE11Ai’s were equally potent and effective with either a 1-hour (Figure S4C, G, K, and O) or 24-hour treatment (Figure S4D, H, L, and P). Further, the effects of the most potent—but not higher—concentrations were reversible following a 5-hour wash out (Figure S4E, I, M, Q and S4F, J, N, and R). To refine our understanding of the time required for PDE11Ai-induced LLPS dispersal, we conducted qualitative live imaging experiments with tadalafil and SMQ-02-57. SMQ-02-57 was chosen based on it being the most potent of our PDE11Ai’s. Both tadalafil (Figure 3A) and SMQ-02-57 (Figure 3B) began dispersing PDE11A4 LLPS in less than 5 minutes. Following washout of tadalafil, PDE11A4 droplets reformed in less than 40 minutes, with the progressive fusion of droplets largely occurring after the first hour (Figure 3C). A follow-up quantitative study using SMQ-02-57 confirmed that levels of PDE11A4 LLPS return to 79% of baseline 1 hour following washout (Figure 3D), with the number of PDE11A4 droplets/LLPS cell doubled at that time point relative to baseline or 5 hours following washout because they have not yet completed progressive fusion (Figure 3E). Such pharmacodynamics are key for considering chronic LLPS-targeted therapy in aging and disease.

**Figure 3.**
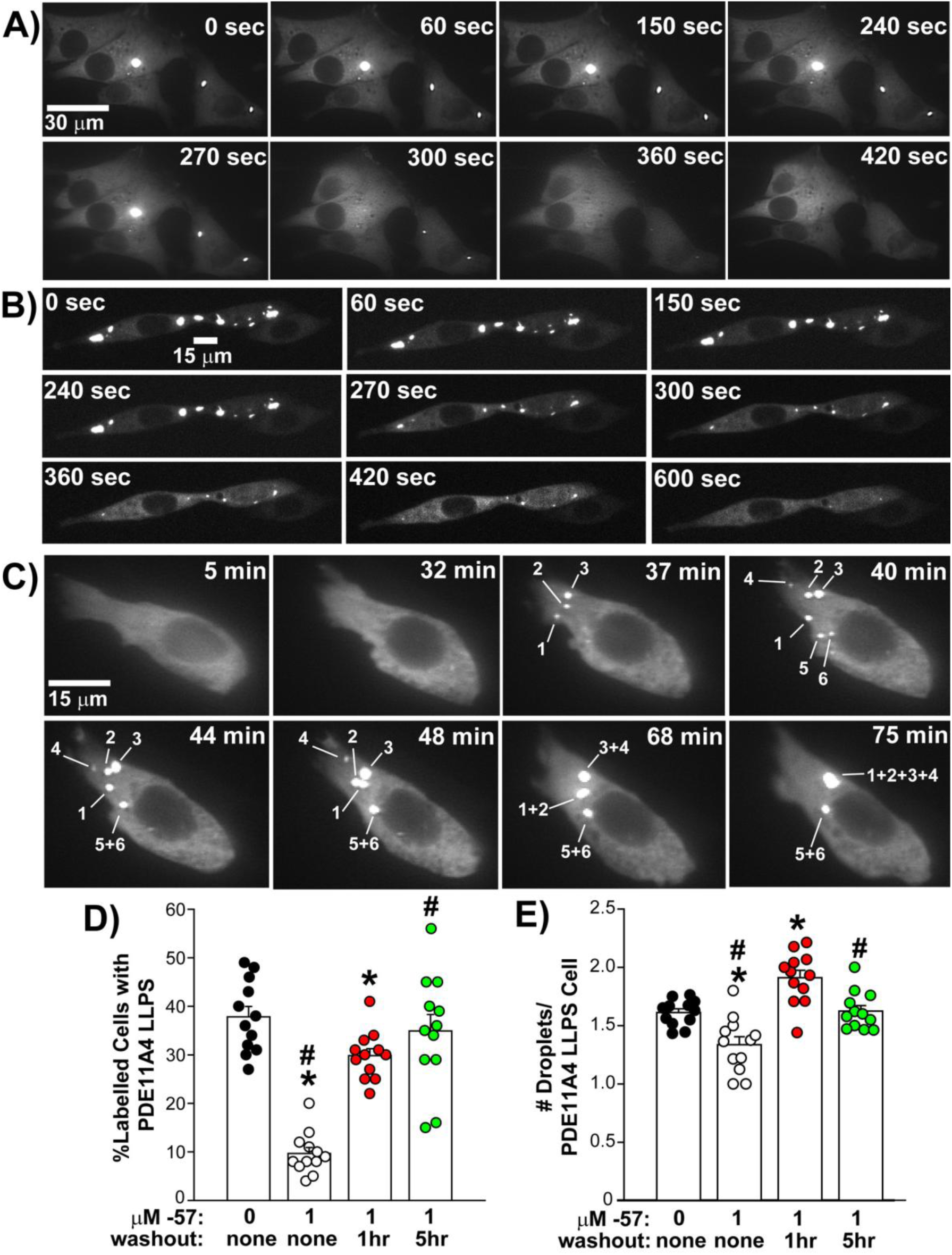
PDE11A4 LLPS reoccurs within 1 hour of washing out a PDE11Ai, with PDE11A4 droplets progressively fusing over time. Time-lapsed imaging of live PDE11A4-transfected HT22 neuronal cells immediately following treatment with **A)** 10 µM tadalafil (representative of 6 cells/3 experiments) or **B)** 10 µM SMQ-02-57 (representative of 16 cells/3 experiments) revealed remixing of PDE11A4 begins within the first 5 minutes of treatment. A follow-up quantitative experiments confirmed that 10 µM tadalafil significantly reduces PDE11A4 LLPS relative to vehicle-treatment levels within 5 minutes (61.7% reduction; n=6/group; t(10)=10.83, P<0.0001). **C)** Time-lapsed imaging of HT22 cells following washout of 10 μM tadalafil (representative of 6 cells over 5 experiments) shows PDE11A4 droplets re-emerge (order indicated by number) and coalesce over time (also see Supplement for video). **D)** Quantification of fixed PDE11A4-transfected HT22 cells that were treated with vehicle, 1 μM SMQ-02-57, 1 μM SMQ-02-57 followed by a 1-hr washout, or 1 μM SMQ-02-57 followed by a 5-hour washout (n=12/group) confirmed that PDE11A4 LLPS begins within the 1st hour of compound removal (equal variance failed; ANOVA on Ranks: H(3)=29.75, P<0.0001; Post hoc: 0 μM vs 1 μM and 1 μM + 1-hour washout P<0.05 for each; 1 μM vs 1 μM + 1-hour washout and 1 μM + 5-hour washout, P<0.05) and that **E)** numerous droplets begin to form but then progressively fuse over time to initially increase and then reduce the total number of PDE11A4 droplets over time (F(3,44)=18.46, P<0.0001; Post hoc: 0 vs. 1 μM and 1 μM + 1-hour washout, P=0.001 for each; Post hoc vs. 1 μM: 1 μM + 1-hour washout P=0.0002, 1 μM + 5-hour washout, P=0.0007). Representative of 2 experiments. *vs. 0 μM, P=0.0016 to <0.0001; #vs. 1 μM treatment, P=0.002-0.0003; @vs. 1hr wash, P=0.0007.

### The PDE11A4 N-terminal regulatory domain contains key sequence features that regulate PDE11A4 LLPS

We next sought to better understand the molecular mechanisms that trigger versus disperse PDE11A4 LLPS. PDE11A4 LLPS requires its N-terminal intrinsically disordered region (IDR), with phosphorylation of S117 and S124 (pS117/pS124) therein increasing PDE11A4 condensation ^10,24^ (see Figure 1D and 1F for location of residues). In contrast, phosphorylation of S162, which falls within the N-terminal regulatory region but outside of the annotated IDR, prevents PDE11A4 LLPS ^24^. We determined if phosphorylation of two additional annotated phosphorylation sites outside of the IDR, S163 and S239, would similarly reduce PDE11A4 LLPS ^33,34^. Since bioinformatic analyses identified LLPS propensity within the GAF-A domain ^10^, we also tested the effect of the PDE11A4-D355A mutation that prevents the functional effects of cGMP allosterically binding to the GAF-A domain ^35^. The phosphomimic mutation S163E reduced PDE11A4 LLPS but the phosphoresistant mutant S163A had no effect (Figure 4A). The phosphomimic S239D also reduced PDE11A4 LLPS while the phosphoresistant S239A promoted it (Figure 4B). This suggests S239, but not S163, is phosphorylated to some extent under basal conditions to limit PDE11A4 LLPS, but there are no phospho-specific antibodies available for these sites to confirm their phosphorylation status. Interestingly, the D355A mutation reduced PDE11A4 LLPS and completely blocked the effects of the aging-like phosphomimic PDE11A4-S117D/S124D (Figure 4C), suggesting this domain critically modulates age-related PDE11A4 LLPS phenotypes. Together, these findings identify novel inroads to therapeutically target PDE11A4 LLPS independently of molecules that also inhibit catalytic activity.

**Figure 4.**
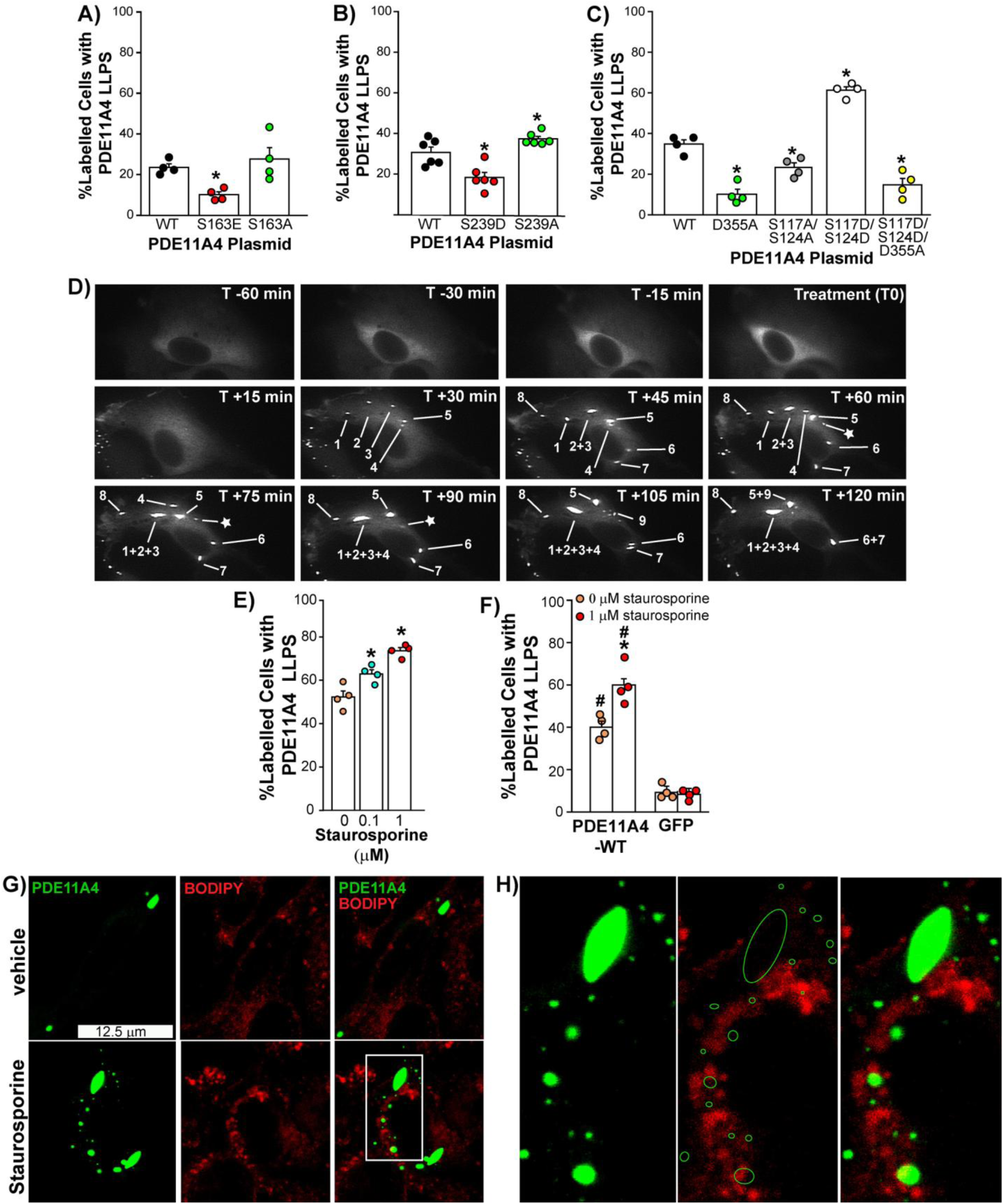
The N-terminal regulatory domain contains key sequence features that regulate PDE11A4 LLPS. Mutations at several residues alters PDE11A4 LLPS including **A)** S163 (n=4 biological replicates/group/experiment; ANOVA: F(2,9)=6.84, P=0.0156; Post hoc vs. WT: S163E P=0.0241, S163A P=0.430), **B)** S239 (n=4 biological replicates/group/experiment; ANOVA: F(2,15)=19.44, P<0.0001; Post hoc vs. WT: S239D P=0.0013, S239A P=0.0493), as well as **C)** D355, S117 and S124 (n=4 biological replicates/group/experiment; ANOVA: F(4,15)=70.92, P<0.0001; Post hoc vs. WT: D355A P=0.0002, S117A/S124A P=0.0042, S117D/S124D P=0.0002, S117D/S124D/D355A P=0.0002; Post hoc vs. D355A: S117A/S124A P=0.0036, S117D/S124D P=0.0001, S117D/S124D/D355A P=0.1679). **D)** Time-lapsed imaging of HT22 cells treated with 0.1 µM of the broad kinase inhibitor staurosporine (n=4 cells/3 experiments). Note the linear form taking shape following fusion of droplets 1-4 (see Supplement for video). **E)** Quantification of droplets in fixed HT22 cells transfected with PDE11A4 and treated with 0-1 µM staurosporine (n=4 biological replicates/group/experiment; ANOVA: F(2,9)=24.59, P=0.0002; Post hoc vs. 0 µM: 0.1µM P=0.0.0069, 1 µM P=0.0003; Post hoc 0.1µM vs. 1 µM P=0.0068). **F)** Quantification of droplets in fixed HT22 cells transfected with either PDE11A4 or GFP alone and treated with 0 or 1 µM staurosporine (n=4 biological replicates/group/experiment; 2-way ANOVA for effect of protein x treatment: F(1,12)=13.25, P=0.0034; Post hoc vehicle vs staurosporine: PDE11A4 P=0.0005, GFP P=0.8107; Post hoc PDE11A4 vs GFP: P=0.0002 for each treatment group). **G)** Representative images of PDE11A4-transfected HT22 cells treated with either vehicle (13 cells/4 experiments) or staurosporine (23 cells/4 experiments) and labeled with the membrane marker BODIPY show no significant colocalization of the two signals (Pearson’s colocalization coefficient: vehicle (r)=-0.006 ±0.005, staurosporine r=-0.003 ±0.004; Manders’ colocalization coefficient M1: vehicle= 0.16 ±0.03, staurosporine=0.11 ±0.02; Manders’ colocalization coefficient M2: vehicle=0.06 ±0.02, staurosporine=0.06 ±0.013). H) Larger magnification of white inset box from panel H with outlines of the PDE11A4 droplet locations overlaid on the BODIPY channel image. A-C and E-F are representative of 2 or more experiments. Post hoc: *vs. WT or 0 µM, P=0.0493-0.0002, #vs. GFP, P=0.0002. Data plotted mean ±SEM.

### Kinases outside of the cAMP/cGMP pathway regulate PDE11A4 LLPS

Given that PDE11A4-p162, -p163, and -p239 outside of the IDR reduce PDE11A4 LLPS, yet activating cAMP/cGMP regulated kinases has no effect on PDE11A4 LLPs ^10^ (replicated in Table S2) and blocking cyclic nucleotide regulated kinases has no effect on PDE11Ai-induced dispersal of LLPS ^10^ (replicated in Table S3), we next determined if broad kinase inhibition beyond the cyclic nucleotide cascade would increase PDE11A4 LLPS. Indeed, the broad kinase inhibitor staurosporine triggered de-mixing of PDE11A4 into spherical droplets that progressively fused over time, a key feature of LLPS (Figure 4D; See Supplement for video of Figure 4D). Further, this fusion led to the formation of a linear condensate (Figure 4D droplets 1-4). This effect of staurosporine on PDE11A4 LLPS occurred in a dose-dependent manner (Figure 4E) and did not extend to GFP alone (Figure 4F). Finally, PDE11A4 spherical droplets in vehicle- and staurosporine-treated cells were membraneless, as shown by a lack of colocalization between our GFP-PDE11A4 signal and the membrane marker BODIPY (Figure 4G). Together these results show that staurosporine increases PDE11A4 LLPS as opposed to enriching its localization within a membrane-bound spherical organelle. Importantly, staurosporine does not increase PDE11A4 LLPS by increasing PDE11A4 protein expression (PDE11A4/Ponceau S relative optical density (r.o.d.) expressed as a fold change of WT: WT 1.00 ± 0.13, 0.1 µM 1.04 ±0.12, 1 µM 1.02 ±0.16; n=9 biological replicates/group; normality failed; ANOVA on Ranks for treatment: H(2)=0.01, P=0.9947. These results support a role for non-cAMP/cGMP kinases in regulating PDE11A4 LLPS and highlight phosphorylation dynamics as a regulatory node.

### PDE11Ai’s reverse PDE11A4 LLPS via a mechanism consistent with allosteric modulation of PDE11A4 homodimerization

As noted above, neither cAMP nor cGMP analogues alter PDE11A4 LLPS and cAMP/cGMP-dependent kinase blockers do not block the ability of PDE11Ai’s to disperse PDE11A4 LLPS ^10^ (see replication experiments in Tables S2-3), suggesting the effect of PDE11Ai’s on PDE11A4 LLPS occurs independently of their catalytic effects. To further investigate the PDE11Ai LLPS-reducing mechanism of action, we examined alternative regulatory pathways. Although mimicking phosphorylation of S163 or S239 reduces basal PDE11A4 LLPS (Figure 4A-B), dispersal of PDE11A4 LLPS by PDE11Ai’s does not require phosphorylation of either S163 (Figure 5A) or S239 (Figure 5B). Instead, native gel Western blots reveal that PDE11Ai’s reduce the abundance of the free PDE11A4 homodimer while shifting PDE11A4 protein to a larger molecular weight complex (Figure 5C-D). The GAF-B domain is required for PDE11A4 homodimerization ^30^, and we previously showed that expression of an isolated PDE11A4-GAF-B domain reduces PDE11A4 homodimerization and mimics the ability of PDE11Ai’s to reduce PDE11A4 LLPS ^23,27^, which we replicate here (Figure 5E). We also previously showed that introducing an A499D mutation within the GAF-B domain stabilizes PDE11A4 homodimerization and increases LLPS ^27^. Therefore, we tested whether A499D would reduce the effectiveness of PDE11Ai’s. As we previously observed, stabilizing homodimerization with the A499D mutation increased PDE11A4 LLPS in vehicle-treated, BC11-38-treated (Figure 5F), and SMQ-02-57-treated cells (Figure 5G). A499D also significantly reduced the effectiveness of PDE11Ai’s across six experiments (Figure 5H; 2 × 100 µM BC11-38, 2 × 1 µM SMQ-02-57, 1 × 10 µM SMQ-02-57, and 1 × 10 µM tadalafil experiments). Further, the effect of A499D was ∼3-fold greater in inhibitor-treated cells than in vehicle-treated cells (Figure 5I). Together, these results support a model in which PDE11Ai binding at the catalytic site allosterically alters PDE11A4 conformation in a manner that reduces homodimerization and, consequently, LLPS.

**Figure 5.**
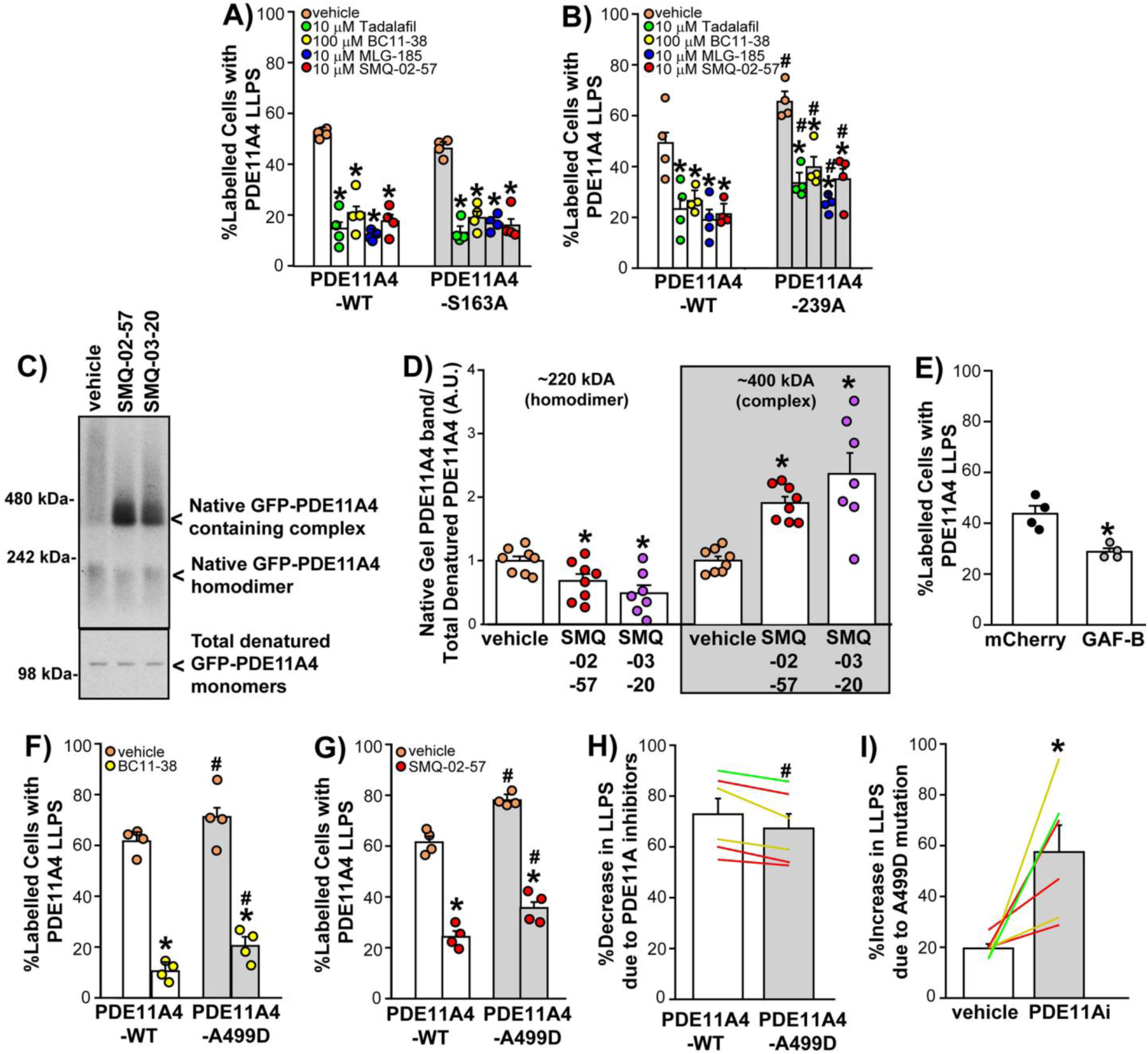
PDE11Ai’s reverse LLPS through mechanisms linked to PDE11A4 homodimerization. Quantification of PDE11A4 LLPS in PDE11A vehicle vs. inhibitor-treated HT22 cells that were transfected with either wild type (WT, n=4/treatment), **A)** phosphoresistant S163A (n=4/treatment; 2-way ANOVA for effect of treatment: F(4,30)=69.4, P<0.0001; effect of treatment x mutant: F(4,30)=1.12, P=0.3667), or **B)** phosphoresistant S239A (n=4/treatment; 2-way ANOVA for effect mutation: F(1,30)=21.48, P<0.0001; effect of treatment: F(4,20)=22.38, P<0.0001; effect of treatment x mutant: F(4,30)=0.44, P=0.7771). **C)** Representative image of Native gel Western (top) and denaturing gel Western blots (bottom) showing PDE11Ai’s reduce the abundance of GFP-PDE11A4 homodimers (migrating ∼220 kDa) and shunt PDE11A4 protein into a macromolecular complex of ∼400 kDa. **D)** Densitometry of PDE11A4 bands from native gel Western blot normalized by the total amount of PDE11A4 protein measured on the denaturing gel Western blot confirms PDE11Ai’s significantly reduced the presence of the 242 kDa PDE11A4 dimer (n=8/treatment; F(2,20)=6.18, P=0.0081; Post hoc vs. vehicle: SMQ-02-57 P=0.0386, SMQ-03-20 P=0.0068) and increased the presence of PDE11A4 in the ∼400 kDa complex (failed equal variance; ANOVA on Ranks: H(2)=13.43, P=0.0012; Post hoc vs vehicle: P<0.05 for each. **E)** Quantification of PDE11A4 LLPS in HT22 cells co-expressing PDE11A4 with either mCherry alone or an mCherry-tagged isolated GAF-B binding domain that reduces PDE11A4 homodimerization (n=4/group; t(6)=4.46, P=0.0043). **F)** Quantification of PDE11A4 LLPS in cells transfected with WT (n=4/treatment) or the homodimerization stabilizing A499D GAF-B domain mutant (n=4/treatment) and treated with either vehicle, BC11-38 (2-Way ANOVA for effect of A499D: F(1,12)=7.29, P=0.0193; effect of treatment: (F(1,12)=199.91, P<0.0001), or **G)** SMQ-02-57 (2-Way ANOVA for effect of A499D: F(1,12)=36.57, P<0.0001; effect of treatment: F(1,12)=298.5, P<0.0001). **H)** Comparison of PDE11Ai anti-LLPS effects between WT and A499D transfected cells (n=6 experiments, (paired t-test: t(5)=4.23, P=0.0082). **I)** Comparison of A499D pro-LLPS effects in vehicle-treated vs. inhibitor-treated cells (paired t-test: t(5)=-3.26, P=0.0223). Post hoc: *vs. vehicle, P=0.0145-0.0001; #vs. WT, mCherry, or 0 μM staurosporine, P=0.0465-0.0001. Data plotted mean ±SEM.

### PDE11Ai’s block the pro-LLPS effects of proteostatic stress, staurosporine, and S117D/S124D

Since age-related LLPS is known to be exacerbated by disease states (c.f., ^8^), we next determined if PDE11Ai’s could block exacerbated PDE11A4 LLPS caused by 1) proteostatic stress, 2) kinase inhibition by staurosporine, or 3) the PDE11A4-pS117/pS124 signature that we previously demonstrated drives age-related condensation of PDE11A4 ^24^. Age-related cognitive decline and dementia are increasingly framed as consequences of a failing proteostasis network—the systems that keep proteins correctly folded, cleared, and non-aggregated ^36^. The secretory pathway is part of the proteostasis network, and the Golgi stress response has been implicated in proteinopathies ^37,38^. The Golgi stress response is triggered when production of secretory and membrane proteins exceeds Golgi processing capacity ^39,40^. Therefore, we modeled secretory-pathway proteostatic stress by overexpressing either TGN38 (a.k.a. TGOLN1), a type I integral Golgi membrane protein that regulates exocytic vesicle formation, or RhoB, a protein that regulates endosomal trafficking and movement ^41–44^. Of direct relevance to the current study, secretory-pathway proteostatic stress can remodel the solubility and condensate behavior of LLPS-prone proteins ^45–47^. Previously we showed that overexpressing TGN38 increases PDE11A4 LLPS in untreated HT22 cells ^21^, and we report here that overexpressing RhoB does the same (n=6/group/x3 experiments; WT + mCherry 52.4 ±1.7 and WT + RhoB 67.5 ±1.9 %labelled cells with PDE11A4 LLPS; t(34)=-6.03, P<0.0001). Here we show that while overexpressing TGN38 increased PDE11A4 LLPS in vehicle-treated HT22 cells, it had no effect in BC11-38-treated cells (Figure 6A). Similarly, overexpressing RhoB significantly increased PDE11A4 LLPS in vehicle-treated cells but not BC11-38-treated HT22 cells (Figure 6B). BC11-38, along with tadalafil, also completely blocked the ability of staurosporine to increase PDE11A4 LLPS (Figure 6C). Finally, multiple PDE11Ai’s across 3 scaffolds blocked the pro-LLPS effect of the aging-mimic S117D/S124D mutant (Figure 6D). This effectiveness of PDE11Ai’s against diverse pro-LLPS triggers underscores the robustness of using small molecules to reduce LLPS and strengthens the argument for therapeutically targeting LLPS in age-related brain dysfunction

**Figure 6.**
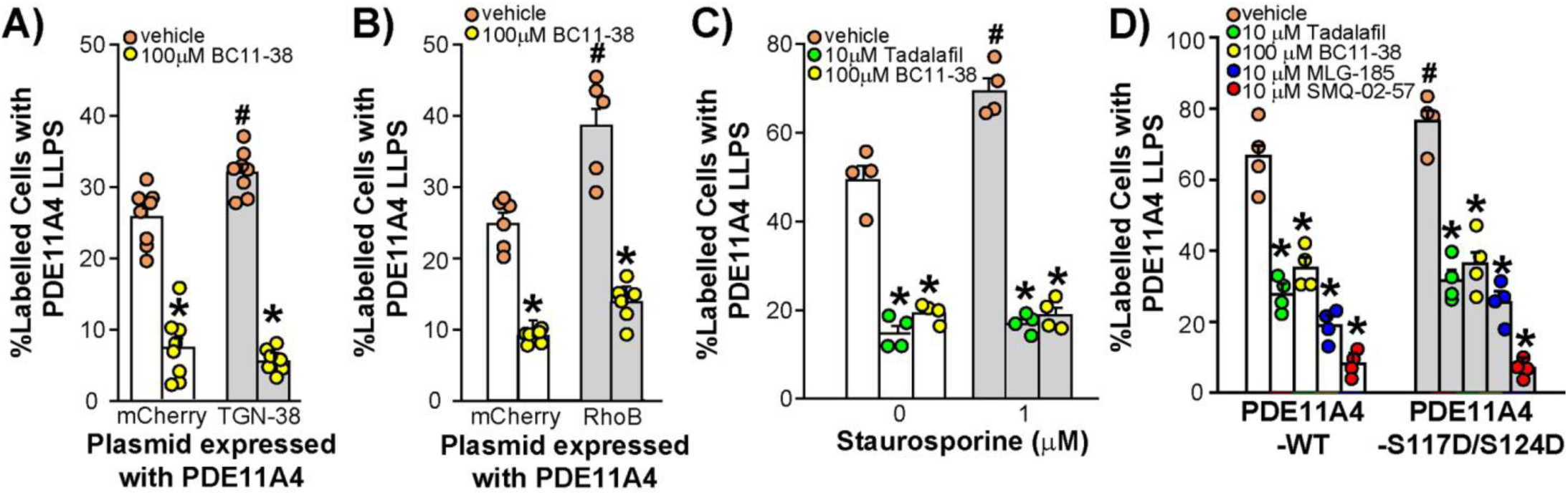
PDE11Ai’s reverse/block pro-LLPS insults. PDE11Ai’s completely reversed or blocked several pro-LLPS mechanisms including **A)** TGN-38 overexpression (a.k.a. TGOLN1; n=8/group; 2-way ANOVA for effect of plasmid x treatment: F(1,28)=11.11, P=0.0024; Post hoc for effect of TGN38 vs mCherry: vehicle P=0.0014, BC11-38 P=0.2645), **B)** RhoB overexpression (n=6/group; 2-way ANOVA for effect of plasmid x treatment: F(1,20)=8.35, P=0.0091; Post hoc for effect of RhoB vs mCherry: vehicle P=0.0002, BC11-38 P=0.0549), **C)** staurosporine treatment (n=4/group; 2-way ANOVA for effect of staurosporine x treatment: F(2,18)=13.36, P=0.0003; Post hoc 0 µM vs 1 µM staurosporine within treatment: vehicle P=0.0002, 10 µM tadalafil P=0.4904, 100 µM BC11-38 P=0.8931), and **D)** the age-related phosphomimic PDE11A4-S117D/S124D mutant (n=4/group; 2-way ANOVA for effect of plasmid x treatment: F(1,30)=4.31, P=0.0465; Post hoc WT vs DDv2: vehicle P=0.0322, tadalafil 0.3831, BC11-38 0.7614, MLG-185 p=0.1451, SMQ-02-57 p=0.7711. Post hoc: *vs. vehicle, P=0.0145-0.0001; #vs. WT, mCherry, or 0 µM staurosporine, P=0.0465-0.0001. Data plotted mean ±SEM.

### Oral dosing of SMQ-03-20 to old NIA C57BL6 mice reverses age-related increases in PDE11A4 ghost axons

Given the promising results in our cell model, we next determined whether oral administration of a brain-penetrant PDE11A4-targeting inhibitor would reverse age-related condensation of PDE11A4-pS117/pS124 in ghost axons, as we previously observed with tadalafil ^10^. We previously demonstrated that SMQ-03-20 is a potent PDE11A4 inhibitor (IC_50_ ∼15 nM) that exhibits only limited inhibition of other PDE families at substantially higher concentrations, and that it shows improved brain penetration relative to tadalafil following oral dosing ^2,48^. At a dose of 30 mg/kg, SMQ-03-20 reaches peak brain exposure at approximately 2 hours (Tmax) and achieves concentrations on the order of ∼5 µM, which is comparable to the EC_50_ for inhibition of PDE11A4 catalytic activity in intact cells (Table S1) and reversal of PDE11A4 LLPS (Figure 1H). Thus, to capture effects on both catalytic activity and subcellular compartmentalization, tissue was harvested 2 hours after oral administration of vehicle (peanut butter alone) or peanut butter pellets containing 30 mg/kg SMQ-03-20. Using this regimen, we previously showed that SMQ-03-20 reduces hypothalamic cAMP-PDE activity in WT but not *Pde11a* knockout mice, supporting PDE11A4-selective target engagement in vivo ^10^. Here, we report that old mice showed significantly higher levels of cAMP-PDE activity relative to young mice in the ventral hippocampus; however, SMQ-03-20 did not significantly inhibit this endpoint (Table S4). This inability to measure a decrease in hippocampal cAMP-PDE activity by SMQ-03-20 was not due to the compound changing PDE11A4 protein levels (Table S4).

Despite the inability to measure a significant inhibition of hippocampal PDE11A4 catalytic activity, SMQ-3-20 did significantly affect other physiological endpoints in the aged hippocampus. Across females and males, SMQ-03-20 significantly reduced age-related increases in PDE11A4 ghost axons by ∼50% in ventral subiculum (Figures 7D, 7I, and S8) and reduced PDE11A4 ghost axons in amygdala to young control levels (Figures 7B, 7I, and S6). SMQ-03-20 also showed a trend toward reducing age-related increases in PDE11A4 ghost axons in vCA1 of old mice (Figures 7C, 7I, and S7) but had no effect on this measure in dorsal CA1 (Figures 7E and S9) or dorsal subiculum (Figures 7F and S10). Neither entorhinal (Figures 7G and S11) nor visual cortex (Figures 7H and S12) demonstrated an age-related increase in PDE11A4 ghost axons, suggesting regional specificity of this age-related phenotype. These data provide additional evidence that the effect of PDE11Ai’s on PDE11A4 LLPS occurs independently of their ability to inhibit PDE11A4 catalytic activity.

**Figure 7.**
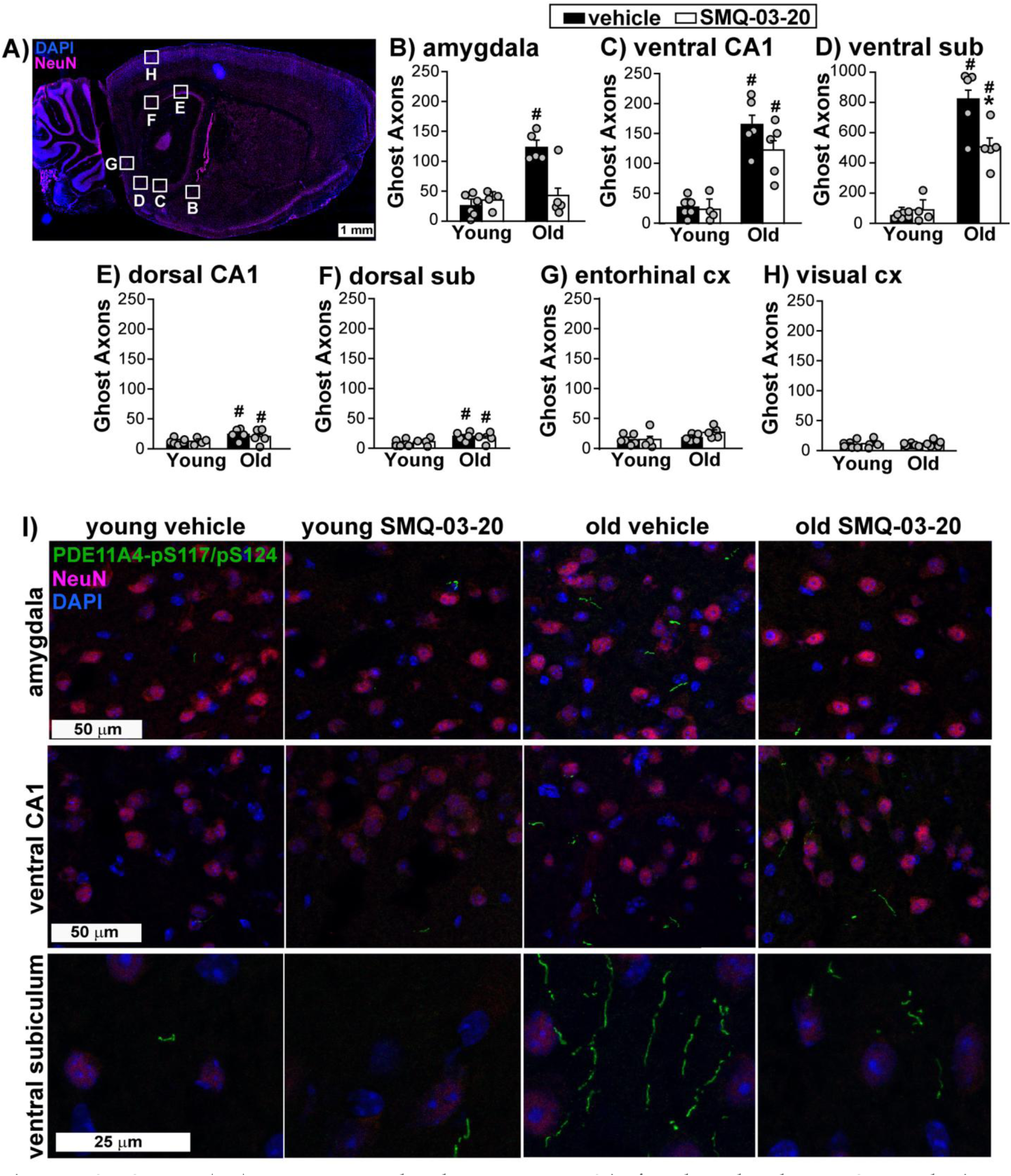
SMQ-03-20 (−20) reverses age-related PDE11A4 LLPS in female and male NIA C57BL6 brains. **A)** Sagittal section from mouse brain outlining brain regions of interest in which the number of PDE11A4 ghost axons was quantified (young vehicle, n=6; young SMQ, n=4; old vehicle and old SMQ, n=5). The letters for each region correspond to the subpanel in which the data are presented. Quantification of PDE11A4 ghost axons in **B)** amygdala (fails normality; ANOVA on Ranks: H(3)=9.57, P=0.0226; Post hoc young vehicle vs old vehicle, P<0.05), **C)** ventral CA1 (2-way ANOVA effect of age: F(1,16)=57.45, P<0.0001; Post hoc: old vehicle vs. old drug, P=0.07), **D)** ventral subiculum (sub; 2-way ANOVA age x treatment: F(1,16)=8.75, P=0.0093; Post hoc vs. old vehicle: young vehicle P=0.0002, old drug P=0.0018), **E)** dorsal CA1 (failed equal variance; Rank Sum Test for effect of age: T(10,10)=75.0, P=0.0254, **F)** dorsal subiculum (2-way ANOVA effect of age: F(1,16)=7.56, P=0.0142), **G)** entorhinal cortex (cx), and **H)** visual cortex. **I)** Fluorescent images of PDE11A4-pS117/pS124 (a marker enriched in PDE11A4 ghost axons), neuron-specific enolase (NeuN, a neuron-specific nuclear marker) and DAPI (a general nuclear marker) from amygdala, vCA1, and ventral subiculum of representative young vehicle-treated, young SMQ-03-20-treated, old vehicle-treated, and SMQ-03-20-treated NIA C57BL6 mice (n=2-3/sex/group). Note that the PDE11A4-pS117/pS124 antibody labels all ghost axons that a pan-PDE11A4 antibody labels (Figure S5) but has better signal:noise ratio and so is used in quantification experiments. See Figures S6-S12 for representative scored images (i.e., PDE11A4 channel only) from each brain region. *vs old vehicle, P=0.0222-0.0018; #vs young, P<0.05-0.0002. Data plotted mean ±SEM.

### Oral dosing of SMQ-03-20 to old NIA C57BL6 mice reverses age-related increases in immunoglobulin light chains

Blood-brain barrier (BBB) integrity declines with aging, particularly the hippocampus^49^. Consistent with this, aged mice demonstrated increased accumulation of immunoglobulins in the hippocampus^50,51^. Thus, we next determined if 30 mg/kg SMQ-3-20 would reverse these age-related increases in hippocampal immunoglobulins. Across female and male NIA C57BL6 mice, SMQ-3-20 reduced age-related increases in ventral hippocampal immunoglobulin light chains by ∼50% (Figure 7M-N). Further, ventral hippocampal immunoglobulin light chains strongly and positively correlated with the number of ghost axons in the ventral subiculum (Figure 8D and H) and ventral CA1 (Figure 8E and I) but not with the number of ghost axons in the dorsal subiculum (Figure 8F and J) or dorsal CA1 (Figure 8G and K), suggesting a localized interaction between ghost axon number and immunoglobulin light chain accumulation. Consistent with the ability of SMQ-03-20 to reduce immunoglobulin light chains in old NIA C57BL/6 mice, *Pde11a* KO mice exhibited only minimal age-related increases in immunoglobulin light chain accumulation (Figure 8C). Importantly, SMQ-03-20 did not decrease immunoglobulin light chain accumulation in the old KO mice. The lack of SMQ-03-20 effects on immunoglobulin light chains in the KO mice proves that SMQ-03-20 reduces immunoglobulin light chain accumulation via its targeting of PDE11A4. Altogether, these findings suggest that reversal of PDE11A4 condensation is associated with the reversal of hallmark pathophysiological changes associated with aging.

**Figure 8.**
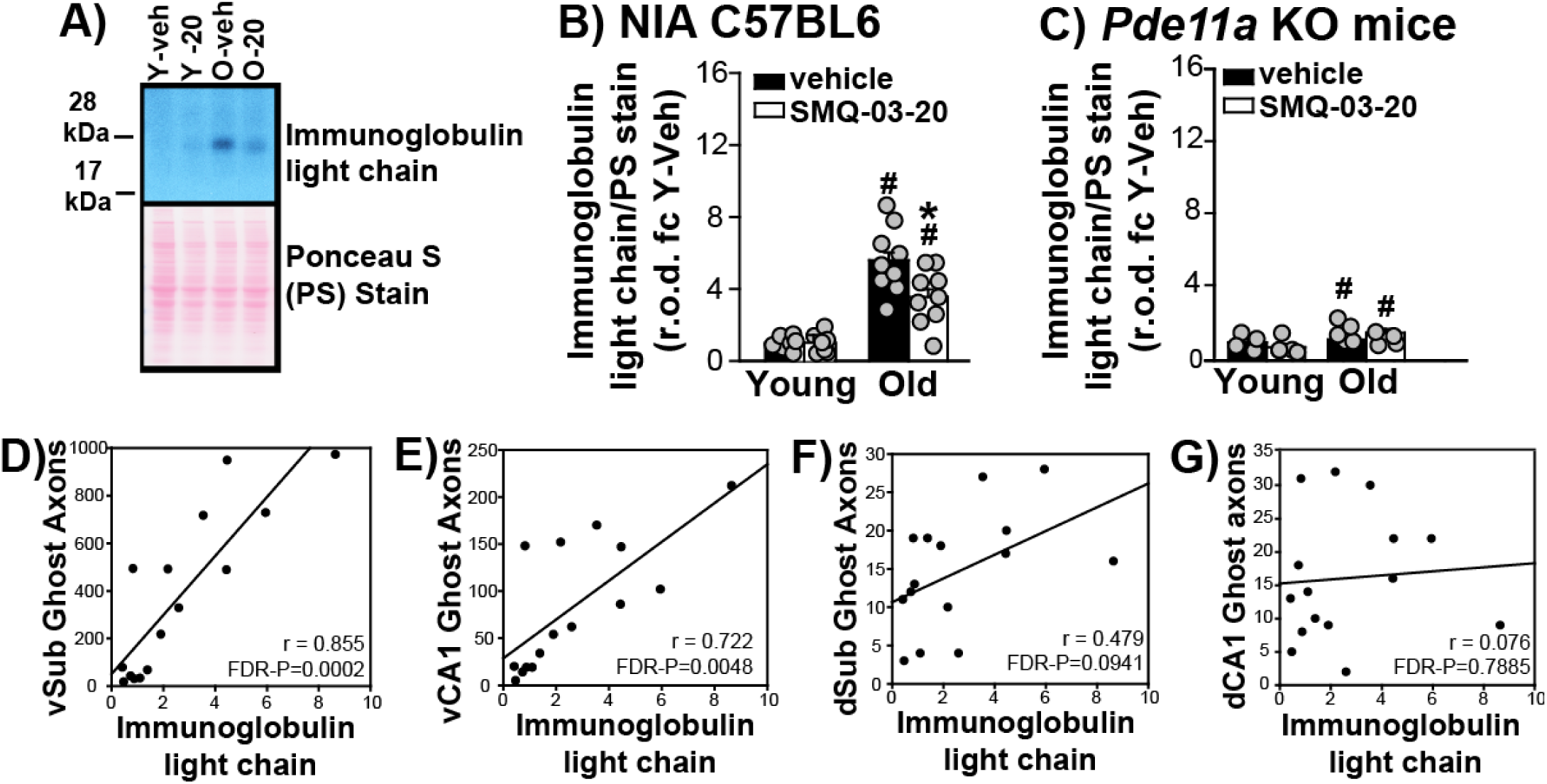
SMQ-03-20 (−20) reverses age-related increases in immunoglobulin light chain accumulation in female and male NIA C57BL6 brain. **A)** Representative Western blot (top) and Ponceau S stained membrane (bottom) from ventral hippocampus of young (Y) and old (O) NIA C57BL6 mice treated with either vehicle (veh) or SMQ-03-20 (−20; n=4-5/sex/group). **B)** Quantification of ventral hippocampal immunoglobulin light chains (failed equal variance; Rank Sum Test (RST) effect of age within vehicle: T(7,9)=28.0, FDR-P=0.004; RST for effect of age within SMQ: T(9,9)=51.0, FDR-P=0.0054; t-test for effect of SMQ within old: t(16)=2.56, FDR-P=0.02813; t-test for SMQ within young: t(14)=-0.1278; FDR-P=0.9001). Relative optical density (r.o.d.) of the immunoglobulin light chains band was normalized by total Ponceau S r.o.d. and expressed as a fold change (fc) of young vehicle. **C)** Quantification of *Pde11a* KO mice ventral hippocampal immunoglobulin light chains (n=2-3/sex/group; 2-way ANOVA for effect of age: F(1,14)=4.97, P=0.0426). **D)** For the 15 subjects with measures for both endpoints, correlations between ventral hippocampal immunoglobulin light chain accumulation and the number of PDE11A4 ghost axons in ventral subiculum (r=0.885, FDR-P=0.0004), **E)** ventral CA1 (vCA1; r=0.722, FDR-P=0.0048), **F)** dorsal subiculum (dSub; r=0.479, FDR-P=0.1129) and **G)** dorsal CA1 (dCA1; r=0.076, FDR-P=0.9012). *vs old vehicle, P=0.028; #vs young, P=0.0426-0.001. Data plotted mean ±SEM.

## DISCUSSION

Using PDE11A4-selective small-molecule inhibitors, we demonstrate that PDE11A4 LLPS and condensates in the aging brain can be reversed, coinciding with reductions in a key marker of BBB dysfunction. We not only identified PDE11A4 LLPS as a regulated, druggable process in the aging brain, but also delineated upstream phosphorylation events, structural motifs, and protein processing pathways that govern its condensation dynamics. Finally, we identified a candidate mechanism by which PDE11Ai’s appear to reverse PDE11A4 LLPS to reveal new therapeutic inroads for targeting this process.

### Reversal of PDE11A LLPS by PDE11A4 small molecule inhibitors

Here, we describe eight PDE11A-targeting small molecules across three chemical scaffolds that consistently and rapidly disperse PDE11A4 condensates (Figure 2), with re-mixing occurring within minutes of exposure (Figure 3A-B). Notably, the six MLG and SMQ inhibitors are highly potent and selective for PDE11A4 ^1,2^ and are predicted to bind PDE11A4 at the catalytic pocket (Figure 1G-H) as previously described for tadalafil ^29–32^. Interestingly, this reversal of LLPS was transient following washout of the most potent concentrations but persisted following washout of higher concentrations (Figure S4). The reason for this is not yet known, but many plausible mechanisms present themselves. There may be both high-affinity and low-affinity binding sites for these PDE11Ai’s, with binding to the high affinity site being reversible and that to the low affinity site being irreversible. The PDE4 family of phosphodiesterases contains the high-affinity rolipram binding site (HARBS) and the low-affinity rolipram binding site ^52^, with rolipram preferentially binding to HARBS and piclamilast—a much more potent PDE4 inhibitor—binding both equally well ^53^. That said, there are no reactive functional groups in any of these PDE11Ai’s, which argues against covalent modification ^1,2^. Alternatively, target-mediated drug disposition (TMDD), where drug target binding alters pharmacokinetics, may play a role ^54,55^. This may be particularly relevant when high compound levels exceed the efflux transporter’s Michaelis constant thereby transitioning the system from efflux dominance to greater intracellular retention ^56–59^. TMDD would then saturate the target and slow inhibitor clearance ^55^. Finally, thermodynamics may differ between low versus high PDE11Ai concentrations, as the formation of drug-target interactions can have unpredictable effects on binding kinetics ^60^. For example, PDE5 transitions from a low-affinity binding state to a high affinity binding state when cGMP allosterically binds its GAF-A domain, thus increasing its sensitivity to inhibition by sildenafil ^61^. If an increased inhibitor concentration stabilizes the drug-target complex without altering the transition state, dissociation may slow ^62^. Determining the mechanism at play here will be of interest to future studies.

PDE11Ai’s also blocked/reversed several pro-condensation triggers, including the age-related pS117/pS124 PDE11A4 signature, proteostatic stressors, and kinase inhibition (Figure 6). Notably, not all LLPS-promoting triggers were equally susceptible: the S239A mutation promoted a drug-resistant form of LLPS (Figure 5B), implying mechanistic heterogeneity in LLPS regulation. The partial specificity of PDE11Ai efficacy may offer opportunities for uniquely targeting distinct PDE11A4 proteinopathies.

### Regulation of PDE11A4 Condensation: A Network of Candidates for Future Therapeutic Targeting

PDE11A4 condensation is orchestrated by a network of phosphorylation events and structural motifs, independent of the enzyme’s canonical catalytic signaling. Within the N-terminal IDR, phosphorylation at S117 and S124 emerges as a critical determinant of age-related PDE11A4 LLPS and ghost axon formation ^24^. Indeed, this IDR is required for PDE11A4 LLPS to proceed ^10^. Phosphorylation at sites outside the IDR (e.g., S162 ^24^; S163, Figure 4A; S239, Figure 4B) also modulates condensate dynamics, but their modification suppresses LLPS. Consistent with this, global kinase inhibition via staurosporine enhances LLPS, producing membraneless droplets that fuse over time (Figure 4D-H) in a reversible manner (Figure 6C). It is then interesting to note that phosphorylation is a key driver of axonal transport deficits associated with Alzheimer’s disease and other dementias ^63–73^ that are phenotypically reminiscent of the PDE11A4 condensation that occurs within ghost axons. Given this phenotypic similarity, it is of interest to future studies to test the effect of PDE11Ai’s on neuronal viability and axonal degeneration, particularly in the context of two rare functional variants in PDE11A4 that have been genetically associated with early-onset Alzheimer’s Disease ^74^. Surprisingly, the phosphorylation events that inhibit PDE11A4 LLPS are not regulated by cAMP/cGMP-regulated kinases ^10^ (Table S2-S3), pointing to a role for non-cyclic nucleotide pathways in regulating PDE11A4 LLPS.

The GAF-A and GAF-B domain within the N-terminal regulatory region of PDE11A4 are also key structural determinants of PDE11A4 LLPS propensity. Disruption of homodimerization via an isolated GAF-B domain reverses PDE11A4 LLPS in vitro and ghost axons in vivo, while stabilization of homodimerization with an A499D mutation promotes both ^23,27,28^. This may not be surprising given that homodimerization is considered a key driver of protein LLPS due to the multivalency of these domains ^75^. A D355A mutation that blocks the functional effects of cGMP allosterically binding the GAF-A domain ^35^ also significantly attenuates LLPS (Figure 4C), aligning with computational predictions of LLPS propensity within this domain ^10^. Together, this body of work demonstrates that both intrinsic sequence features and extrinsic signaling converge on a tightly regulated LLPS program for PDE11A4, thus, opening new therapeutic avenues for targeting this process.

### PDE11Ai’s Mechanism of Action Is Consistent With Disrupting Homodimerization Rather Than Enzyme Inhibition

Contrary to expectation, PDE11Ai’s reverse LLPS independently of catalytic inhibition. More specifically, we previously demonstrated and replicate here (Table S3) that blocking cAMP/cGMP kinases had no impact on inhibitor-induced reversal of PDE11A4 LLPS ^10^. We also previously showed ^10^ and replicate here (Table S2) that neither cAMP nor cGMP analogues mimicked the LLPS-reversing effect of PDE11Ai’s. Similarly, inhibition of other PDE families had no effect on PDE11A4 LLPS (Figure 2L-M), supporting the specificity of PDE11A4-targeting molecules. PDE11Ai’s also acted independently of changing PDE11A4 protein expression (Table S1) or increasing S163 or S239 phosphorylation (Figure 5A-B). Instead, PDE11Ai’s reduced the abundance of PDE11A4 homodimers, shifting PDE11A4 to higher molecular weight complexes (Figure 5C-D). Further, disrupting PDE11A4 homodimerization mimicked the ability of PDE11Ai’s to reduce LLPS (Figure 5E), while strengthening homodimerization with an A499D mutation increased LLPS (Figures F-H) and attenuated the effect of PDE11Ai’s on LLPS (Figure 5H). Although the identity of the higher molecular weight complex has yet to be identified, our results are consistent with disrupted homodimerization being a key mechanism by which PDE11Ai’s reduce LLPS. That said, we cannot rule out the possibility that additional mechanisms— such as changes in post-translational modifications or interactions with scaffolding proteins —also contribute.

The crystal structure of PDE11A has yet to be solved; however, biochemical and modeling data indicate that tadalafil binds PDE11A in the catalytic pocket and that a single hydrogen bond to Gln-869 is critical for high-affinity binding ^29–32^. Our modeling here suggests that our PDE11A-selective inhibitors also bind the catalytic domain and interact with Gln-869 (Figure 1G-H). Thus, our working hypothesis is that binding of these small molecules to the PDE11A4 catalytic pocket induces a conformational change that allosterically disrupts homodimerization (Figure S13). This hypothesis is supported by several observations: (i) in other PDE families, catalytic-site ligand binding induces conformational states that allosterically couple to distal domains, including dimer interfaces ^76–84^; (ii) PDE11A’s oligomerization is sensitive to N-terminal/catalytic-region organization ^30,85^; and (iii) 1,6-hexanediol—a general multivalency disruptor—disperses PDE11A4 LLPS ^10^. Direct testing of this hypothesis will require additional structural and biophysical studies, which will be facilitated by future resolution of the PDE11A4 structure.

### Oral SMQ-03-20 Reverses Age-related Increases in Ghost Axons and Immunoglobulin Light Chains

In vitro, PDE11A4 condensates largely appear as discrete spherical droplets, but short linear structures can be observed (e.g., Figure 4D “1+2+3+4”; also see Figures 4D-E and S9 in ^24^). Further, while ghost axons in vivo appear filamentous at low magnification, higher-resolution imaging reveals ghost axons are often composed of closely spaced spherical or ovoid puncta arranged along a linear trajectory (Figure 7I and S5-S12). Although direct in vivo imaging of PDE11A4 LLPS was beyond the scope of the present manuscript, we suggest that PDE11A4 spherical droplets observed in vitro and PDE11A4 ghost axons observed in vivo represent different stages of the same underlying process. In the staurosporine time-lapse experiment (Figure 4D), PDE11A4 first formed numerous spherical droplets within ∼30 minutes, followed by droplet fusion at ∼45 minutes and the emergence of elongated linear assemblies by ∼75 minutes (also see supplemental video). The apparent differences between cellular and in vivo morphologies therefore likely reflect differences in timescale and protein accumulation. Our cell model only expressed PDE11A4 for several hours, whereas the age-related accumulation in vivo spans months allowing for continued growth and packing of condensates. Importantly, PDE11A4 ghost axons are reversible in vivo ^10,23^ (Figures 7A and 7D), which supports a dynamic, regulated assembly consistent with a condensate state as opposed to stable filamentous aggregates. Thus, the cellular and in vivo phenotypes are best interpreted as temporally distinct manifestations of PDE11A4 LLPS.

Oral administration of the highly potent and PDE11A-selective small molecule SMQ-03-20 significantly reduced PDE11A4 ghost axons of aged NIA C57BL/6 mice, eliminating condensation in the amygdala and reducing it by ∼50% in ventral subiculum (Figures 7B, 7D). Some regions (e.g., dorsal subiculum) were unaffected by the compound, possibly due to regional variability in drug exposure (e.g., ^86,87^) or regional differences in scaffolding partners mediating homo-vs. heterotypic LLPS. The dispersing effects of SMQ-03-20 in the ventral hippocampus occurred without measurable changes in cyclic nucleotide hydrolysis in this brain region. We cannot rule out the possibility that localized reductions in PDE11A4 activity may be occurring within the ventral hippocampus that fall below our threshold for detection. That said, we were able to measure a significant inhibition of cAMP-PDE activity in the hypothalamus of these same SMQ-03-20-treated mice ^2^ and in the ventral hippocampus of two different *Pde11a* KO mouse lines ^20,28^. The reversal of PDE11A4 LLPS in ventral hippocampus in absence of detectable changes in bulk ventral hippocampal PDE activity is consistent with our previous findings showing that inhibition of cAMP/cGMP-dependent kinases fails to block PDE11Ai reversal of LLPS and cAMP/cGMP analogues fail to mimic the effect of PDE11Ai’s on LLPS ^10^ (Tables S2-S3). LLPS is regarded as an emergent phenomenon with a characteristic switch-like response, with weakening of multivalent interactions dissolving droplets in an effectively all-or-none manner ^88–91^. In contrast, catalytic inhibition generally follows a continuous, hyperbolic dose–response, with functional inhibition requiring higher occupancy to produce graded changes in bulk activity ^92,93^. Thus, we suggest our observations are most consistent with a mechanist model in which modest PDE11A4 inhibitor binding collapses LLPS in a thresholded, switch-like process without measurably changing bulk catalytic activity (a graded process).

Notably, the reversal of ventral hippocampal age-related PDE11A4 ghost axons by SMQ-03-20 coincided with a 50% reduction in immunoglobulin light chain protein, that strongly correlated with ghost axon counts in ventral subiculum and ventral CA1 but not dorsal subiculum or dorsal CA1 (Figure 8). The accumulation of immunoglobulins in the aging hippocampus has been associated with BBB dysfunction^49–51^. Thus, it is of great interest to identify the underlying mechanism by which neuronal PDE11A4 regulates neuroimmune and neurovascular processes, particularly in the context of aging. Critically, the effect of SMQ-03-20 on immunoglobulin light chain accumulation was absent in *Pde11a* knockout mice (Figure 8C), suggesting the PDE11Ai effects were mediated via PDE11A4 specifically. Aberrant phase separation has recently been implicated in BBB dysfunction^94^, but our work here provides the first evidence linking PDE11A4 LLPS to age-related neuroinflammatory markers.

Although direct, real-time characterization of condensate dynamics in vivo was beyond the scope of the present study, several lines of evidence support the physiological relevance of PDE11A4 compartmentalization in vivo. Age-related increases in ventral hippocampal PDE11A4 protein occur selectively within the membrane-associated fraction, with no corresponding increase in cytosolic or nuclear fractions ^24^. Moreover, disrupting PDE11A4 homodimerization using an isolated GAF-B peptide selectively reduces this membrane-associated PDE11A4 pool, prevents PDE11A4 LLPS in cell models, reverses ghost axon accumulation in the old mouse brain, and rescues memory deficits in old mice ^23,27^. In contrast, social isolation of young adult mice selectively reduces ventral hippocampal PDE11A4 protein expression in the membrane-associated fraction, which is sufficient to drive subsequent changes in social behaviors ^28,95,96^. Together, these findings support the existence of a regulated, optimally compartmentalized state of PDE11A4 in vivo that is mechanistically linked to functional outcomes and is consistent with the concentration-dependent condensation behavior observed in cells.

### Study Limitations

While the present study provides mechanistic insight into molecular mechanisms that regulate PDE11A4 LLPS and demonstrates promising effects of PDE11Ai’s both in vitro and in vivo, several limitations of the study should be acknowledged. First, neuronal viability, axonal maintenance, synaptic functional readouts nor behavioral outcomes were measured. Further, while we demonstrated that reversal of PDE11A4 LLPS in vivo corresponds with reversal of age-related increases in immunoglobulin light chains, we did not assess BBB function directly. These constraints should be considered when interpreting our findings. Despite these limitations, we believe this work is timely and provides a compelling rationale to continue exploring the therapeutic implications of reversing protein-specific LLPS in the context of aging and neurodegenerative disease.

## METHODS

### Compounds

Tadalafil (Tocris, 6311), BC11-38 (medchem express, Hy108618), rolipram (Sigma, R6520), papaverine (Sigma, P3510), and staurosporine (medchem express, Hy15141) were obtained from commercial vendors. MLG-199, MLG-185, MLG-122, SMQ-02-57, SMQ-03-30, and SMQ-03-20 were synthesized by the Rotella lab as previously described ^1,2^.

### Computational method

AlphaFold3 ^97^ was used to model the structure of PDE11A4 based on the amino acid sequence from UniProt entry Q9HCR9-1. The predicted catalytic domain exhibited very high pLDDT scores, indicating strong confidence in the structural model. Site Identification by Ligand Competitive Saturation (SILCS) ^98^ was then applied to characterize functional group binding patterns within the catalytic domain of the AlphaFold3 derived structure. SILCS simulations were conducted to generate functional group free energy maps (FragMaps) according to the previously established protocol ^98^. Subsequently, Monte Carlo (MC) sampling using these FragMaps (SILCS-MC) ^99^ was performed to globally dock the compounds of interest within the SILCS-Hotspots framework ^100^ and predict their favorable binding modes to PDE11A4. The docking pose with the most favorable ligand grid free energy (LGFE) score was selected as the most probable binding mode for further analysis.

### Plasmid generation

Plasmids were generated as previously described ^27^. Briefly, Genscript (Piscataway, NJ) generated constructs expressing either EmGFP alone containing an A206Y mutation to prevent EmGFP dimerization ^101^ or the mouse *Pde11a* (NM_001081033) or human *Pde11a4* (NM_016953.4) sequence fused at the N-terminal with EmGFP. Note that mouse *Pde11a* is ∼95% homologous and the same length as human *Pde11a4* and so the protein is referred to herein as PDE11A4 for clarity. These constructs were initially generated on a pUC57 backbone and then subcloned into a pcDNA3.1+ mammalian expression vector (Life Technologies; Waltham, MA). RFP-tagged RhoB (Cat# RC100059) and RFP-tagged TGOLN/TGN38 plasmids (Cat# RC100061) were obtained from OriGene.

### Cell culture and transfection

As described above, PDE11A4 expression in brain is enriched in the hippocampus ^20,25,102^. Therefore, we use the HT22 hippocampal cell line (sex undefined) for most *in vitro* investigations ^24^. As previously described ^1,24^, cells were maintained in T-75 flasks in Dulbecco’s Modified Eagle Medium (DMEM) with sodium pyruvate (GIBCO, Gaithersburg, MD or Corning, Manassas, VA), 1% Penicillin/Streptomycin (P/S); GE Healthcare Life Sciences; Logan, UT), and 10% fetal bovine serum (FBS; Atlanta Biologicals), with incubators set to 37°C/5% CO_2_. Cells were passaged at ∼70% confluency using TrypLE Express (GIBCO; Gaithersburg, MD). The day before transfection, cells in DMEM+FBS+P/S were plated in 60 mm dishes, 24-well plates, or coverslips depending on the experiment. The day of transfection, the media was replaced with Optimem (GIBCO) and cells were transfected using 1 microliter Lipofectamine 2000 (Invitrogen; Carlsbad, CA) per mL of Optimem, as per the manufacturer’s protocol. In experiments testing only a single plasmid, 0.375 µg of cDNA was used for transfections. For studies employing more than 1 plasmid, pilot studies were conducted to empirically determine the amount of plasmid needed per plasmid in order to yield equivalent levels of total protein expression across plasmids. ∼19 hours post-transfection, the Optimem/Lipofectamine solution was replaced with DMEM+FBS+P/S. Cells continued growing for five hours in the supplemented media and then followed 1 of 4 courses. They were either 1) stained with 2µM BODIPY-568 in (Invitrogen, #D3835) 1x phosphate-buffered saline (PBS 10x Powder Concentrate, Fisher BioReagents) for 15 minutes and then fixed in 4% paraformaldehyde and mounted using DAPI Fluoromount-G mounting media (Southern Biotech), 2) directly fixed in 4% paraformaldehyde (Sigma-Aldrich) in 1x PBS and then stored in 1x PBS, 3) pharmacologically treated for 1 hour or 24 hours and then fixed as above and stored in 1X PBS, or 4) treated for 1 hour or 24 hours, then switched to compound-free DMEM+FBS+P/S for 5 hours and then fixed and stored in 1X PBS.

Images of cells labeled with Bodipy-568 were captured using the Leica Model DM6 CFS confocal microscope in the University of Maryland School of Medicine (UMSOM) Department of Neurobiology Imaging Core equipped with the Leica TCS SP8 laser system, Leica STP8000 control panel, Lumencor Sola Light Engine epifluorescent lamp, and Leica CTR6 power source using a 63X/1.40 Oil CS2 ∞/0.17/OFN25/E HC PL APO objective. Static images for quantification of PDE11A4 compartmentalization into spherical clusters were collected from cells stored in 1x PBS using a Nikon Eclipse TE2000-E Inverted via a 10x/0.40 CS2 ∞/0.17/OFN25/A objective equipped with Photometrics CoolSNAP cf camera and CoolLED pE-300lite LED illuminator. Representative images for each well were captured using MetaVue v6.2r6 software and saved as jpeg files. This Nikon microscope is located in the UMSOM Department of Neurobiology Imaging Core. Over the course of experiments, cells were sporadically tested for yeast, fungal, and bacterial infections (Invitrogen; Cat#:C7028), with negative results always obtained.

### Quantification of PDE11A4 LLPS

Because transient transfections can yield different levels of transfection efficiencies across experiments, LLPS data are expressed as the percentage of labeled cells exhibiting PDE11A4 LLPS. LLPS-positive cells contain bright, punctate structures that are readily identifiable and scored with high consistency. In contrast, cells exhibiting only diffuse cytosolic PDE11A4 labeling are inherently lower in contrast and may vary in detectability due to differences in expression level or the contrast of individual rater set ups. As such, variation in the total number of labeled cells (denominator) can arise between raters due to differences in identifying these lower-intensity cytosolic cells. Thus, the baseline %LLPS does vary between raters/experiments. To ensure consistency, all images within an experiment were quantified by an experimenter blinded to treatment using the same computer, monitor settings, zoom level, room lighting conditions, and position in the room, as previously described ^10,21,23^. Experimenters were trained using reference datasets containing constructs known to increase or decrease PDE11A4 LLPS to ensure a given rater’s equipment enabled a sufficient window of measurement to detect both increases and decreases from baseline. All statistical comparisons were performed within an experiment, where scoring conditions were held constant.

### Time-lapsed imaging

Cells were plated on Mattek poly-D-lysine coated dishes (Mattek, P35GC1.514C) but otherwise cultured and transfected as described above. Cells treated with staurosporine were then imaged using an Olympus VivaView LCV110 Fluorescence Incubator Microscope equipped with a 40x: UlanSApo 40x/0.95 infinity symbol/0.11-0.23/FN26.5 objective and a Hamatsu Orca-R2 C10600 camera operated by MetaMorph software (Molecular Devices, San Jose, CA) set to a 20x digital magnification. For staurosporine experiments, HT22 cells were imaged in drug-free supplemented media for 1 hour to find cells that did not spontaneously develop PDE11A4 LLPS (note: only about 40% of cells develop PDE11A4 LLPS at this time point). Imaging then continued for 2 hours after adding the staurosporine. Cells treated with SMQ-02-57 were plated in 24-well plates and imaged using a Nikon Eclipse TE2000-E Inverted via a 10x/0.40 CS2 ∞/0.17/OFN25/A objective equipped with Photometrics CoolSNAP cf camera and CoolLED pE-300lite LED illuminator. Images were captured using MetaVue v6.2r6 software and saved as jpeg files and then assembled into a video using ImageJ. Cells were allowed to form PDE11A4 droplets and then were imaged before and in 30-second increments following treatment with 10 µM SMQ-02-57.

### PDE activity assay

As previously described ^10^, cells were treated for 1 hour, after which the media was removed, the cells were harvested in PDE assay buffer (20mM Tris-HCL and 10mM MgCl_2_) and homogenized using a tissue sonicator (output control: 7.5, duty cycle: 70, continuous). Samples were then held at 4 °C until processing. Total protein levels were quantified using the DC Protein Assay Kit (Bio-Rad, Hercules, CA) according to the manufacturer’s directions. 3 µg of each sample was then processed for cGMP- and/or cAMP-PDE activity using a radiotracer assay based on ^103^, with some adjustments ^1,2^. Briefly, samples were incubated with 35000-45000 disintegrations/minute of[^3^H]-cAMP or[^3^H]-cGMP for 10 minutes. The reaction was then quenched with 0.1M HCl and neutralized using 0.1M Tris. Snake venom was then added to the sample and incubated for 10 minutes at 37 °C. Samples were then run down DEAE A-25 Sephadex columns previously equilibrated in high salt buffer (20 mM Tris-HCl, 0.1% sodium azide, and 0.5M NaCl) and low salt buffer (20 mM Tris-HCl and 0.1% sodium azide). After washing the columns four times with 0.5 mL of low salt buffer, the eluate was mixed with 4 mL of scintillation cocktail, and then CPMs were read on a Beckman-Coulter liquid scintillation counter. 2 reactions not containing any sample lysate were also taken through the assay to assess background, which was subtracted from the sample CPMs. Data was expressed as CPMs/ug protein.

### Western Blotting

As previously reported ^24^, cells were sonicated in ice cold lysis buffer (20 mM Tris-HCl, pH 7.5; 2 mM MgCl_2_; Thermo Pierce Scientific phosphatase tablet #A32959 and protease inhibitor 3 #P0044) or in PDE assay buffer as described above. Each sample destined for native blots was then treated with universal nuclease to degrade DNA (0.25 U nuclease/10 µl of sample). Protein concentrations were determined by DC Protein Assay kit (Bio-Rad; Hercules, CA, USA) according to manufacturer protocol, and were subsequently equalized across samples. Samples were stored at −80°C until further processing. For western blotting, 10µg of total protein was loaded onto 4-12% NuPAGE (i.e., denaturing) or NativePAGE Bis-Tris mini protein gels (Invitrogen, Waltham MA) and electrophoresed for one at 180 volts (denaturing) or a total of 3.5 hours at 150 volts (native). GFP-transfected cell samples were included on all PDE11A4 blots as a negative control. Protein was transferred onto a 0.45µm nitrocellulose (denaturing) or PVDF (native) membrane (Amersham, #10600008) for two hours at 100 mA. Nitrocellulose membranes were then washed twice in Tris-buffered saline with 0.1% tween20 (TBS-T) before staining with Ponceau S to determine total protein loading on denaturing gels. Note, Ponceau S was chosen over a housekeeping gene as a loading control based on the best-practice statement of the Journal of Biological Chemistry ^104^. Images of the stained membranes were collected to later quantify the optical density of the total protein stain (i.e., spanning ∼200kDa to 10kDa), and then the membranes were rinsed in TBS-T to remove the stain. PVDF membranes were first fixed in 8% acetic acid and then washed in TBS-T. Blots to be probed with our custom PDE11A4 antibody (chicken polyclonal; Aves, #1-8113a; 1:10,000) were blocked in 5% milk while those to be probed with GFP (rabbit polyclonal; Santa Cruz, #sc8334; 1:2000), Histone H3 (Millipore, #05-928, 1:20,000), pAKT (Cell Signalling, #9271S, 1:1000), Synaptophysin (BD

Biosciences, #611880, 1:200,000), IL-1β (Invitrogen, #M421B; 1:80,000), and IL-6 (Life Technologies, #ARC0962; 1:300,000) were blocked in Superblock Blocking Buffer (ThermoFisher, Cat#37515), each with 0.1% Tween 20. Primary antibodies were incubated overnight at 4°C. The next day, membranes were washed 4 × 10 minutes with TBS-T and then incubated for 1 hour at room temperature with a secondary antibody (anti-chicken: Jackson Immunoresearch, 103-035-155 at 1:40 000; anti-rabbit: Jackson Immunoresearch, 111-035-144 at 1:10 000; anti-mouse: Jackson Immunoresearch, 115-035-146 at 1:10,000). Subsequently, membranes were washed 3 x 15 minutes in TBS-T. Finally, the membranes for IL-1β or IL-6 were immersed in WesternSure Premium Chemiluminescent substrate (LI COR, #926-9500) while all other membranes were immersed in SuperSignal West Pico Chemiluminescent Substrate (ThermoScientific, Waltham MA) as per manufacturer’s directions. Following this step, membranes were wrapped in clear a plastic sheet protector and exposed to film. Multiple film exposures were taken to ensure signals were within the linear range, and Ponceau S stain and western blot optical densities were quantified using Image J. To account for membrane-membrane variances in film exposure, antibody saturation, chemiluminescence reaction, etc. between blots, Western blot data were normalized to a control condition (e.g. GFP-PDE11A4 + vehicle) on each blot, as previously described (e.g., ^25,105,106^).

Please note that, due to a 100-fold dilution miscalculation, the IL-6 and IL-1β primary antibodies were used at concentrations insufficient to detect their intended targets (Figure S14A-B). The resulting blots showed indistinguishable banding patterns, which were subsequently reproduced in a control blot incubated with anti-mouse secondary antibody in the absence of primary antibody (Figure S14C). These findings demonstrated that the bands originally attributed to IL-6 and IL-1β instead reflected endogenous mouse immunoglobulin detected by the anti-mouse secondary antibody, with the band at the relevant molecular weight corresponding to immunoglobulin light chains. Unfortunately, insufficient tissue remained from most animals in the study, so we were only able to test n=4 subjects/group on the secondary-only control blot, which was not sufficiently powered for formal analyses. Accordingly, all group comparisons and correlation analyses were performed using the original full-cohort “IL-6” (main text Figure 8) or “IL-1β” blots (Figure S14E-J).

### In vivo study

All in vivo experimental protocols were approved by the Institutional Animal Care and Use Committee at the University of Maryland, Baltimore with treatment and care conducted as per the National Institutes of Health Guide for the Care and Use of Laboratory Animals (Pub 85-23, revised 1996). All methods are reported in accordance with ARRIVE guidelines. As previously described ^2,10^, young C57BL6 mice were imported from the National Institute of Aging (NIA) colony and were then bred onsite at UMSOM. *Pde11a* KO mice on a mixed C57BL6 background (99% C57BL/6J and C57BL/6NCrl, 1% 129P2/OlaHsd as per MiniMuGA background panel; Transnetyx, Cordova, TN) were also bred onsite at UMSOM. All mice were group-housed 4-5/cage, with young mice tested at ∼3 months old and old mice tested at ∼18 months old. Equal numbers of female and male mice were used and no animals were excluded from analyses. Within each cage of mice, ½ were designated to receive vehicle and ½ to receive drug. To randomly assign each mouse a treatment within a cage, identification numbers for mice were sorted in ascending order. Treatment group was then assigned in alternating order down the list of IDs, with half of cages starting with vehicle and half of cages starting with drug (i.e., vehicle, drug, vehicle, drug vs. drug, vehicle, drug, vehicle). Post hoc analyses confirmed there was no difference in body weights between vehicle and drug-assigned mice.

As previously described ^2^, we used peanut butter as a vehicle for oral delivery of SMQ-03-20. Briefly, Jif brand creamy peanut butter was melted to a liquid state in a sterile beaker by stirring it on a warming plate. The liquid peanut butter was either used plain or had a body weight-appropriate amount of SMQ-03-20 added to yield a 30 mg/kg dose to mice. The plain peanut butter was then slowly poured into molds using a 3 mL syringe without a needle to avoid bubbles or voids. The rectangular cavities of the mold hold 100 µL of liquid and a sterile razor blade was used to scrape off any excess peanut butter that escaped the cavity. The SMQ-03-20-laden peanut butter was poured similarly into separate SMQ-03-20-designated molds. Molds were then placed onto dry ice for ∼20 minutes to freeze and then were stored in an ultralow freezer until time of use. On the day of testing, pellets were removed from the freezer and immediately placed on dry ice where they remained until the precise time at which a mouse was dosed. Mice were food restricted for 1 night and the next morning (Day 1), all mice were transported to a quiet (∼50-52 dB), brightly lit room (∼700-800 lux) designated for *in vivo* studies. Mice were then singly housed in clean plastic cages with no bedding, allowed to habituate for 1 hour, and then provided a plain peanut butter pellet for habituation. Typically, mice took ∼10-60 minutes to eat the first pellet. At this point, ad lib food was returned to the home cage. On day 2, mice were again habituated to a plain peanut butter pellet, taking ∼7-8 minutes to consume the pellet. On day 3, half of the mice were provided a plain peanut butter pellet (vehicle) and the other half were given a peanut butter pellet containing a 30 mg/kg dose of SMQ-03-20. Generally, mice ate their pellets in ∼2-3 minutes.

2 hours after consuming their pellet, mice were moved to a second room and were killed by manual cervical dislocation (without anesthesia or other chemical agent) by an experimenter blind to treatment. As previously described ^24^, tissue was harvested fresh on wet ice, flash frozen in 2-methylbutane sitting on dry ice, placed directly on the dry ice to allow for evaporation of the 2-methylbutane. ½ brains were then stored long-term at −80° C until being processed via immunofluorescence.

### Immunofluorescence (IF)

Fresh-harvested, flash-frozen brains were embedded in matrix, cryosectioned in the sagittal plane at 20 µM, and thaw-mounted onto a +/+ slide. As previously described ^24,102^, frozen tissue was fixed using 4% paraformaldehyde in 1X PBS for 10 minutes. After fixation, the tissue was washed 3 × 10 minutes in 1X PBS with bovine serum albumin and triton X-100 (PBT). For quantification, overnight incubation was conducted at 4°C using an antibody cocktail in PBT that included 2 antibodies that detect PDE11A4-pS117/pS124, a posttranslational modification found almost exclusively in clustered PDE11A4 ^24^ (Fabgennix PPD11A-140AP at 1:1000 and Fabgennix PPD11A-150AP at 1:500; rabbit). For figure images, slides were incubated overnight with this cocktail plus an antibody for NeuN (Millipore Sigma MAB377 at 1:1000; mouse) or this cocktail plus our custom pan-PDE11A4 antibody (Aves #1 at 1:10,000; chicken). The following day, sections were washed 4x in PBT and incubated for 90 min in secondary antibody (Jackson Immunoresearch #711-545-152 Alexafluor 488 Anti-Rabbit at 1:1000 for cocktail; Jackson Immunoresearch #711-585-152 Alexafluor 594 AffiniPure Anti-Rabbit at 1:1000 for cocktail only when co-labeling with #1; Jackson Immunoresearch #703-545-155 Alexafluor 488 AffiniPure Donkey Anti-Chicken at 1:1000 for #1; Jackson Immunoresearch #711-585-150 Alexafluor 594 AffiniPure Anti-Mouse at 1:1000 for NeuN). The secondary antibody was then washed off with PBT in 3X10 min washes. After rinsing the slides 10X in 1X PBS, slides were then treated with TrueBlack Lipofuscin Autofluorescence Quencher (Biotium, catalog # 23007). After, excess TrueBlack was rinsed off with agitation 3X in 1X PBS. Once dry, slides were mounted with DAPI fluoromount (Southern Biotech, #0100-20). Slides were kept covered and refrigerated until imaged using a Leica Model DM6 CFS microscope equipped with the Leica TCS SP8 laser system, Leica STP8000 control panel, Lumencor Sola Light Engine epifluorescent lamp, and Leica CTR6 power source. Representative images were captured with the Leica Application Suite X software using the HC PL APO 20x/0.74 IMM CORR CS2 objective with max projection images from Z-stacks saved as TIFF files.

### Counting ghost axons

Endogenous PDE11A4 ghost axons in brain were quantified by an experimenter blind to treatment. PDE11A4 “ghost axons” are filamentous accumulations of PDE11A4 that can appear continuous or as a trail of dots. Ghost axons were quantified using the consistent application of classification standards with consideration to the continuity of the strand, quality of brightness, and proximity with other ghost axons.

Analyses of images were counter-balanced such that one image per group was analyzed before moving onto the second image from a group. Non-specific or dull structures were omitted, while non-continuous strands of dots required further consideration in determining whether one or multiple ghost axons were present. Data accuracy was maintained by conducting each analysis under consistent room lighting, monitor settings, and time of day across all samples. Images were quantified using the “Analyze” function in the “ImageJ” application. The magnification was set to 75% and remained constant in all images. Data was gathered using the “Multi-Point” tool which incrementally recorded the number of ghost axons upon each click. The total number of ghost axons was then entered into an .xls spreadsheet for subsequent analysis.

### Data analyses

All between-group analyses were performed using Sigmaplot v11.2. Age and/or treatment effects were analyzed by One-way or Two-way ANOVA (F) when normality as per the Shapiro-Wilks test and equal variance as per the Levene’s test passed. When these assumptions were not met, then an ANOVA on ranks (H) or Mann Whitney Rank Sum Test (T) was used, with FDR-correction applied to P values to account for multiple testing. Following a significant main effect or interaction, *post hoc* tests were conducted using Student Newman-Keuls method and significance was defined as P<0.05. Please note that Sigmaplot provides exact P-values for *post hoc* tests following parametric ANOVAs, but only yes or no to “P<0.05” for *post hoc* tests following a significant non-parametric ANOVA. Data are graphed as mean ± standard error of the mean (SEM). Source data, unadjusted/uncropped images, and videos time-lapse images can be found in Supplemental Materials.

## Supporting information

Amurrio et al., uncropped images

Ammurio et al., supplemental figures and tables

Amurrio et al. Figure 1F_PDE11A4 homodimer rotating

Amurrio et al., Figure 3C tadalafil washout LLPS

Amurrio et al, Figure 4D staurosporine LLPS

## SUPPLEMENTAL MATERIALS

- Supplemental Tables and Figures: Amurrio et al Supplemental Materials.doc
- Videos: Figure 1F_rotating PDE11A4 homodimer.mp4, Figure4D_PDE11A4_LLPS_after_staurosporine.avi, and Figure3C_PDE11A4 LLPS after Tadalafil washout.avi
- Source data: Amurrio et al Source Data.pdf
- Original images of blots: Amurrio et al unprocessed blot images.pdf

## DECLARATIONS

### Ethics approval and consent to participate

In vivo studies were reviewed and approved by the Institutional Animal Care and Use Committee at the University of Maryland, Baltimore with treatment and care conducted as per the National Institutes of Health Guide for the Care and Use of Laboratory Animals (Pub 85-23, revised 1996).

### Consent for publication

Not applicable.

### Availability of data and materials

The datasets supporting the conclusions of this article are included as a supplemental file and are archived here: https://data.mendeley.com/datasets/g3xt59g92y/1

### Authors’ contributions

Investigation, writing parts of the original draft and reviewing the entire manuscript: all authors. Visualization: EA, MD, HG, WY, and MPK. Resources and Funding acquisition: MPK and DPR. Conceptualization, methodology, formal data analyses, and data curation: MPK.

## Acknowledgements

Authors would like to thank Latarsha Porcher, Madison Goodwin, Adrita Sanyal, Sophie Bruckmeier, Asmaa Hijazi, and Om Bhardwaj for technical assistance. The authors would also like to acknowledge Charles S Hoffman, Jeremy Eberhard, Dennis Colussi, John Gordon, and Wayne Childers for generating in vitro screening data reported elsewhere ^1,2^ that contributed to decisions about which compounds we tested herein.

ScholarGPT and Consensus were used to conduct select literature searches, improve clarity, and reduce word count.

## Data are patent pending

◦ 2025 US Non-Provisional Patent Application no. 18/998,725. Inhibitors of PDE11A4 and methods of using the same. [extension of US Provisional Patents 63/370,546 and 63/393187]
◦ 2025 US Non-Provisional Patent Application no. 18/998,724. Inhibitors of PDE11A4 and methods of using the same. [extension of US Provisional Patent 63/393,167]
◦ 2025 European patent application for Entry into the European National Phase no. EP23847591.7. [based on International Patent WO 2024/26500]
◦ 2024 International Patent WO 2024/026500 A2 (pending), Inhibitors of PDE11A4 and methods of using the same.
◦ 2024 US Provisional Patent Application no. 63/692,455. Inhibitors of PDE11A4 and methods of using the same.
◦ 2022 US Provisional Patent US 63/370,546. Inhibitors of PDE11A4 and methods of using the same.

## LIST OF ABBREVIATIONS

BBB: blood-brain barrier
cAMP: cyclic adenosine monophosphate
cGMP: cyclic guanosine monophosphate
cx: cortex
dCA1: dorsal CA1
DMEM: Dulbecco’s Modified Eagle Medium
dSub: dorsal subiculum
GFP: emerald-green fluorescent protein
FBS: fetal bovine serum
FDR: false-discovery rate
FragMaps: functional group free energy maps
GAF: cGMP-regulated phosphodiesterases (PDEs), Anabaena adenylyl cyclases, and Escherichia coli transcription factor FhlA
HARBS: high affinity rolipram binding site
IDR: intrinsically disordered region
LGFE: ligand grid free energy
LLPS: liquid–liquid phase separation
KO: knockout
M1 or M2: Manders’ colocalization coefficient
MC: Monte Carlo sampling
NeuN: neuron-specific enolase
NIA: National Institute of Aging
PBS: Phosphate buffered saline
PBT: PBS with bovine serum albumin and triton X-100
P/S: penicillin/streptomycin
PDE11A4: phosphodiesterase 11A4
PDE11Ai’s: PDE11A inhibitors
pLDDT: predicted Local Distance Difference Test
r.o.d.: relative optical density
RFP: red fluorescent protein
RST: Rank Sum Test
SILCS: Site Identification by Ligand Competitive Saturation
TBS-T: Tris-buffered saline with 0.1% tween20
TMDD: target-mediated drug disposition
UMSOM: University of Maryland School of Medicine
vCA1: ventral CA1
vSub: ventral subiculum
WT: wild type

