## Supplementary material for "Molecular mechanisms regulating PDE11A4 age-related liquid-liquid phase separation and its reversal by selective PDE11A4-targeted small molecules in cells and brain": Amurrio et al., uncropped images

**Figure 2A)**

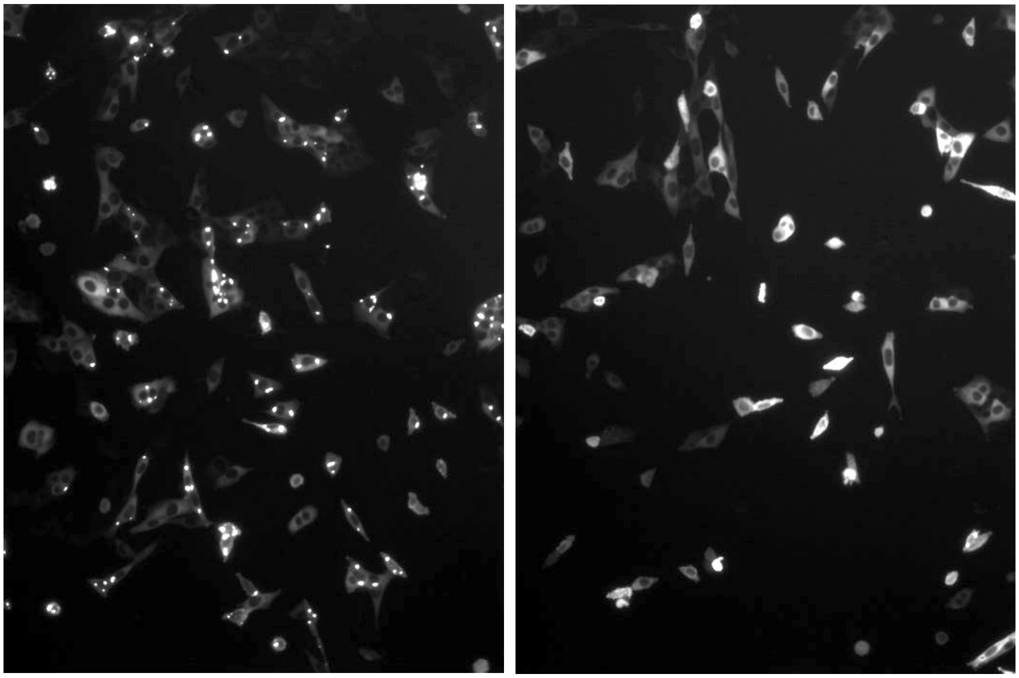

**Figure 2E)**

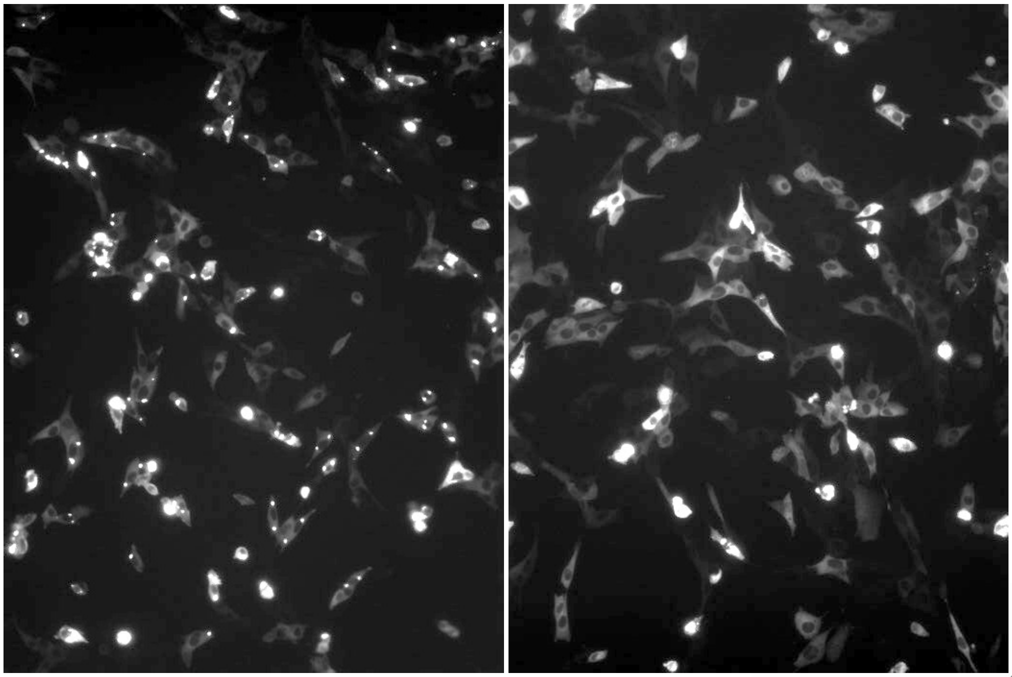

**Figure 2I)**

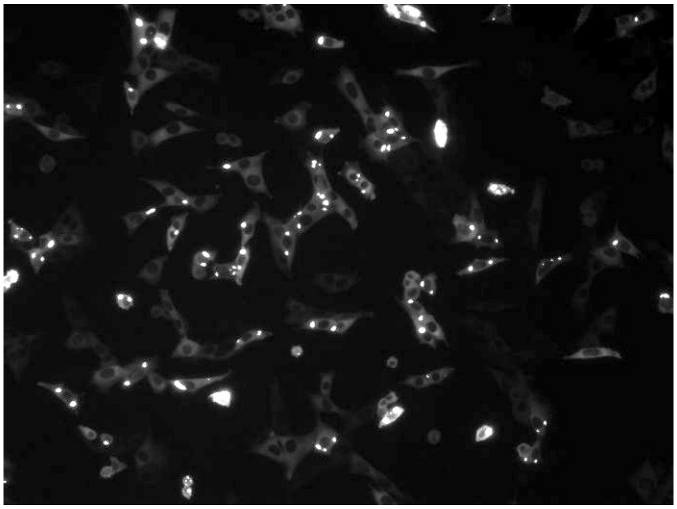

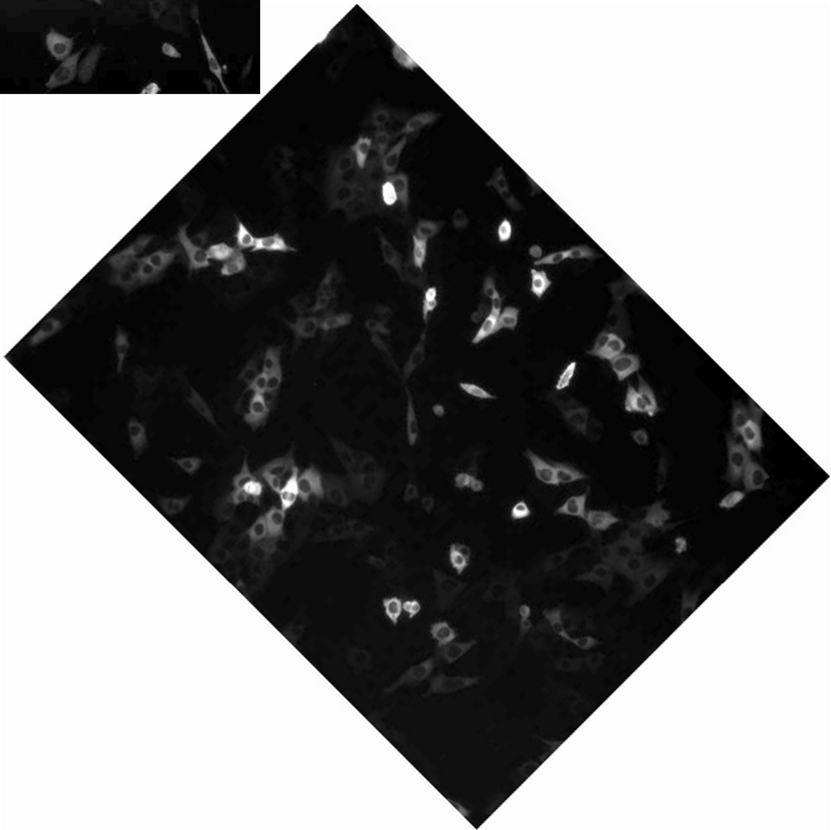

### Figure 5C

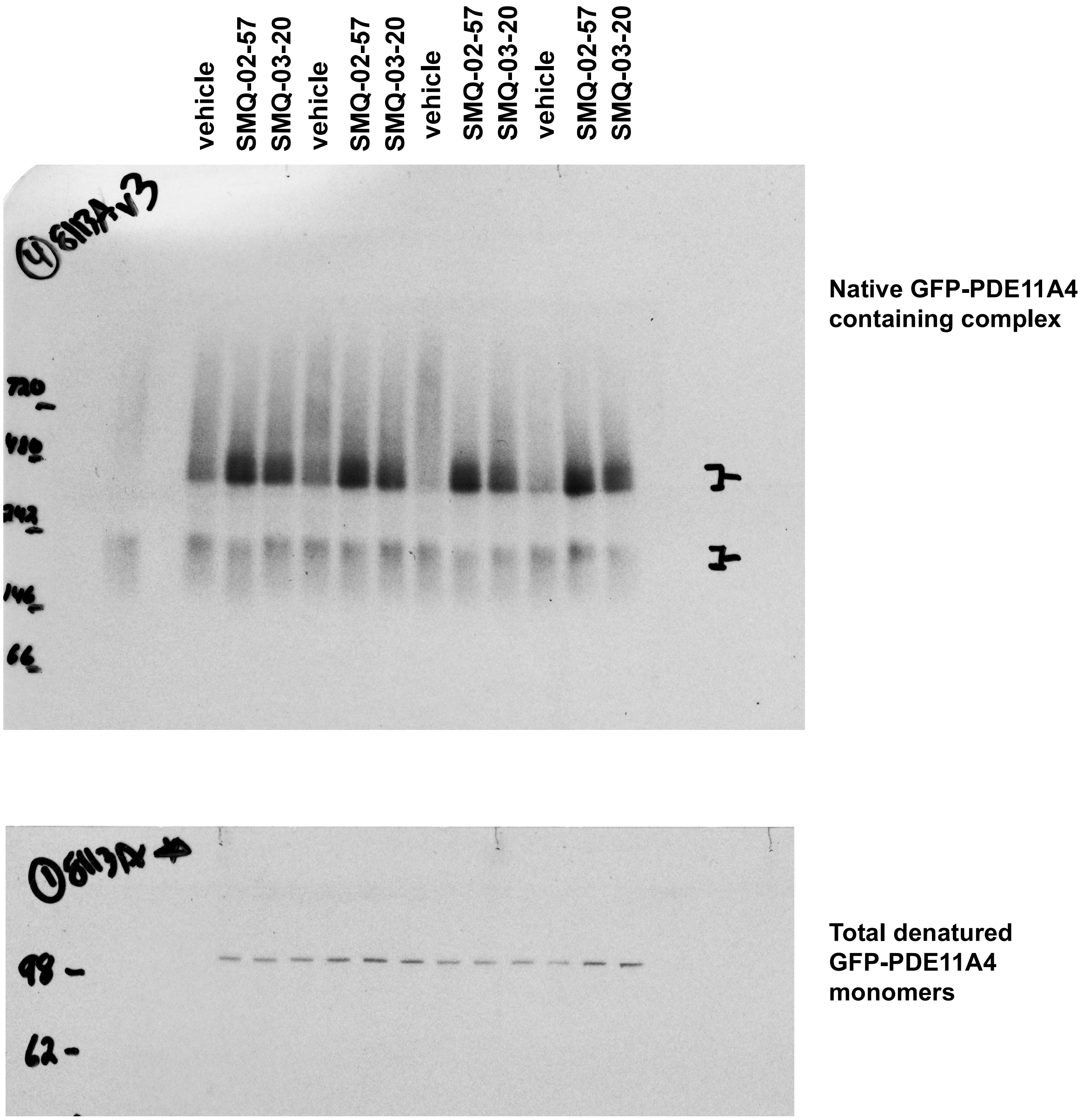
vehicle
SMQ-02-57
SMQ-03-20
vehicle
SMQ-02-57
SMQ-03-20
vehicle
SMQ-02-57
SMQ-03-20
vehicle
SMQ-02-57
SMQ-03-20
720
480
242
146
66
Native GFP-PDE11A4 containing complex
98
62
Total denatured GFP-PDE11A4 monomers

#### Figure 8A

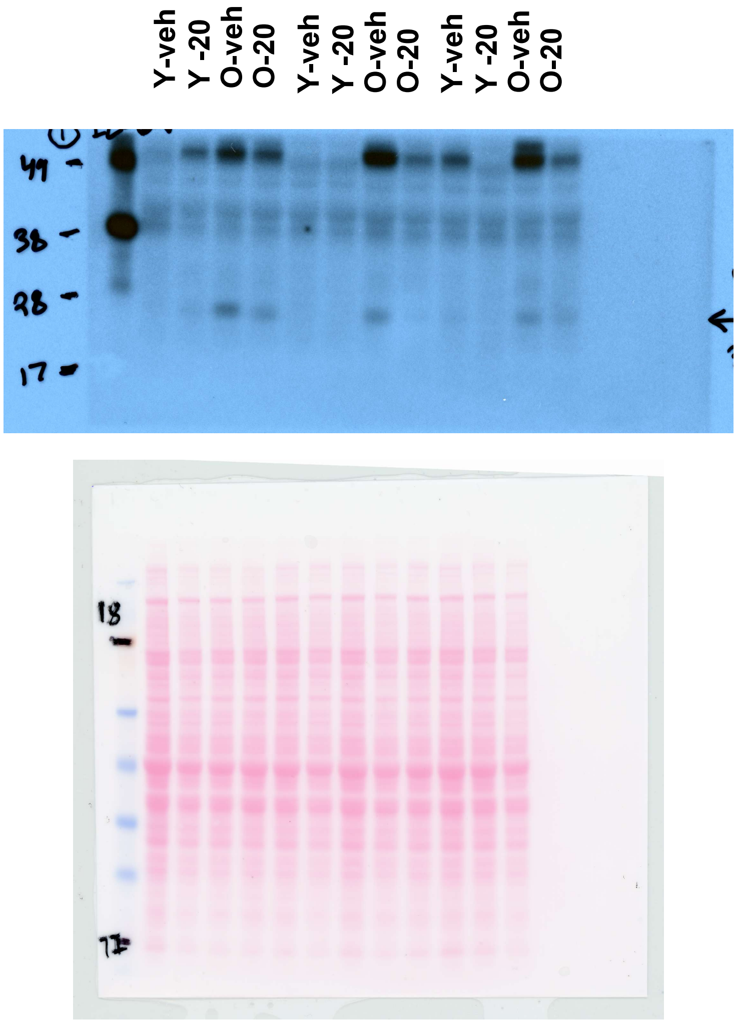
Y-veh
Y -20
O-veh
O-20
Y-veh
Y -20
O-veh
O-20
Y-veh
Y -20
O-veh
O-20
49
38
28
17
18
17

FigureS1

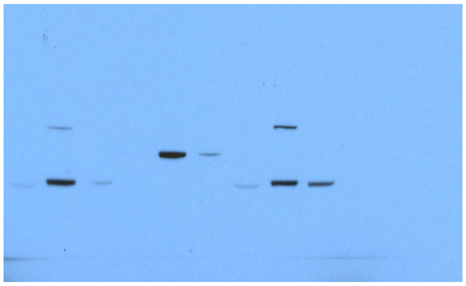

BLOTS:

PDE11A4

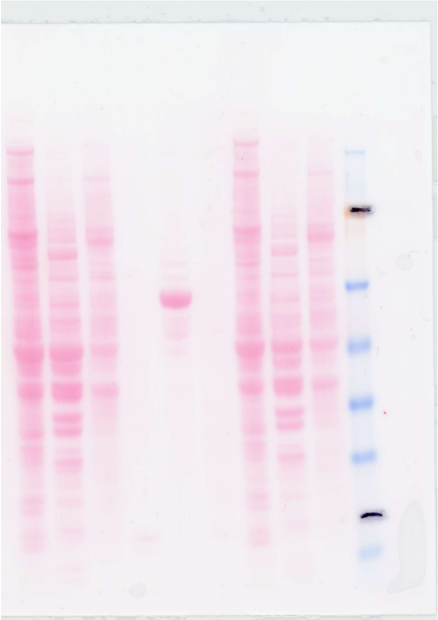

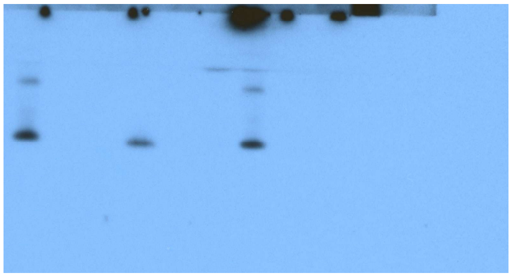

H3

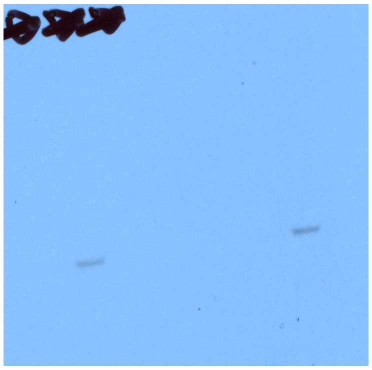

AKT, short exposure for tissue

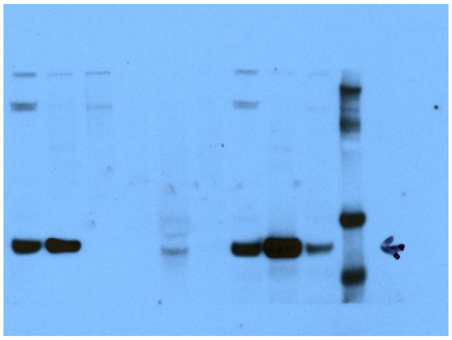

AKT, long exposure for cells

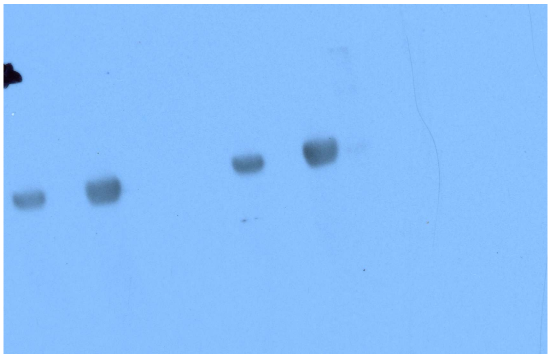

synaptophysin as membrane marker for tissue

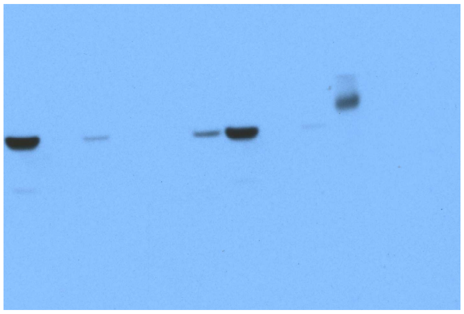

VDAC1 for membrane marker for cells

**Figure S2**

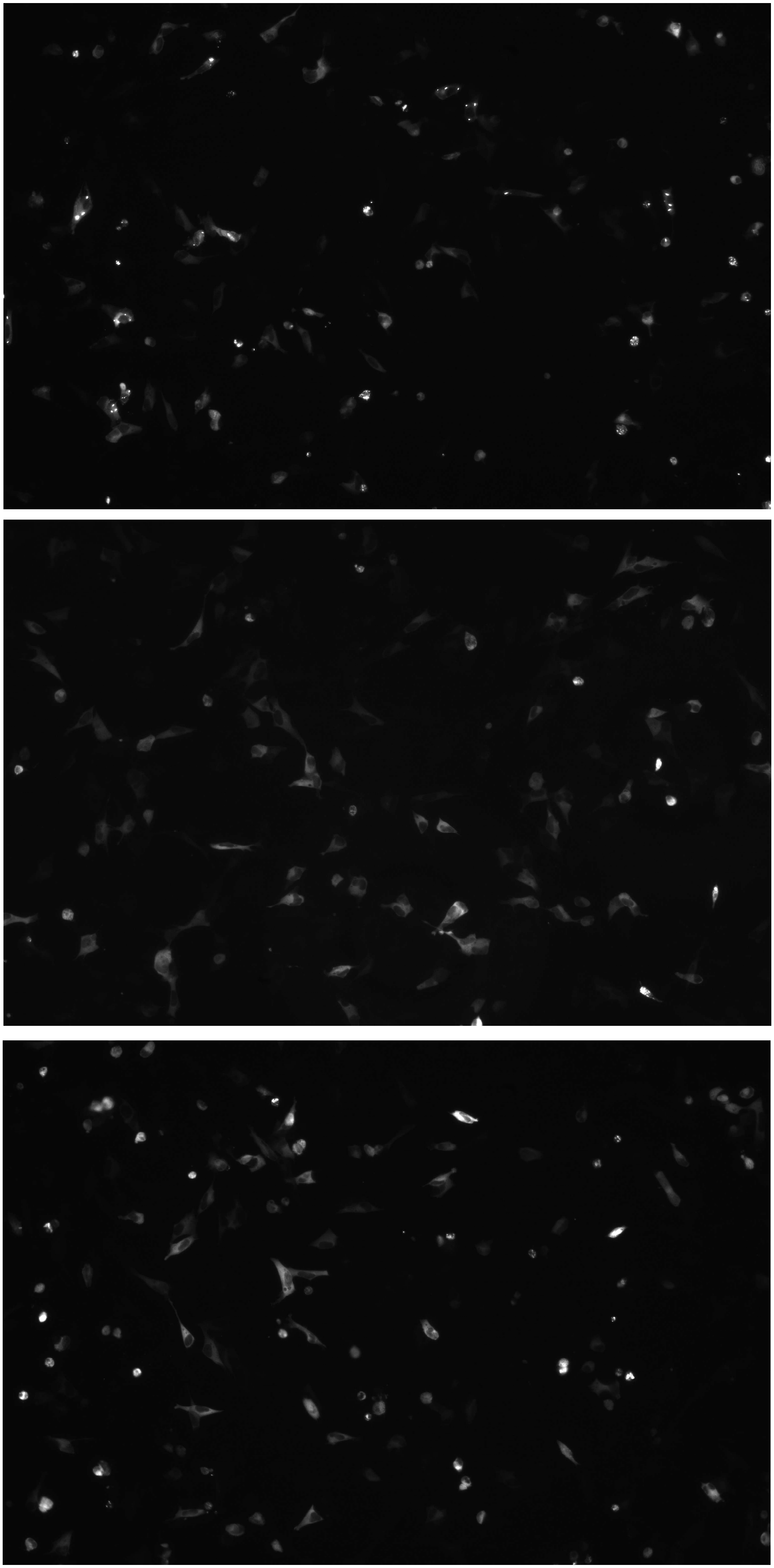

**Figure S4**

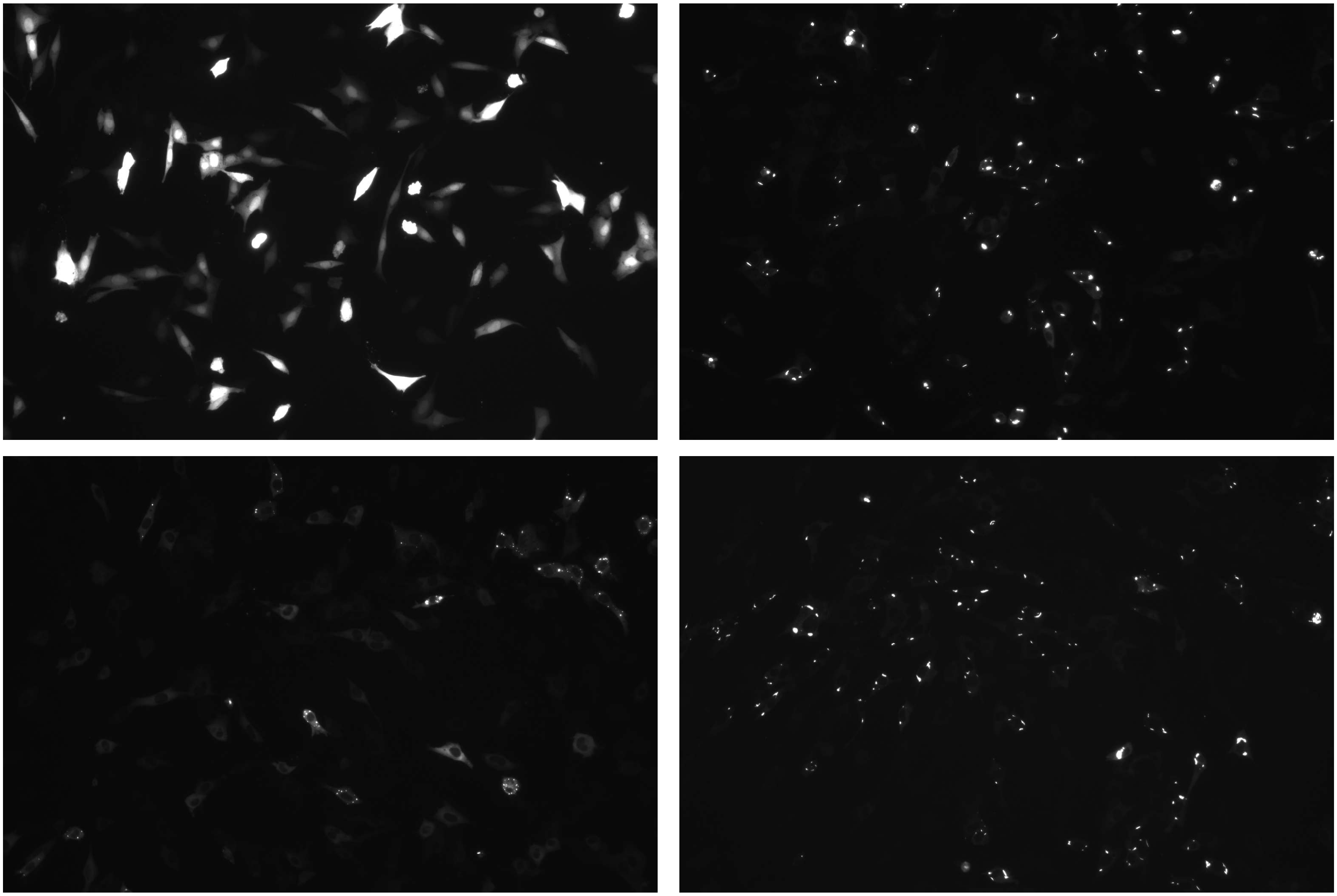
