## Supplementary material for "Molecular mechanisms regulating PDE11A4 age-related liquid-liquid phase separation and its reversal by selective PDE11A4-targeted small molecules in cells and brain": Ammurio et al., supplemental figures and tables

**Table S1. PDE11A inhibitors identified in screens against human PDE11A4 also inhibit GFP-tagged mouse PDE11A4 cAMP- and cGMP-catalytic activity in mouse HT22 hippocampal neuronal cells without significantly decreasing total PDE11A4 protein expression levels.**

| | GFP + 0 μM | WT + 0 μM | WT + 0.1 μM | WT + 1 μM | WT + 10 μM | WT + 100 μM | statistical results |
|---|---|---|---|---|---|---|---|
| **tadalafil** | | | | | | | |
| cAMP-PDE | **0.03±0.01** | 1.0±0.03 | 0.99±0.12 | 1.02±0.05 | 0.66±.04 | **0.15±0.01** | H(5)=19.01, P=0.0019 |
| cGMP-PDE | **0.01±0.00** | 1.0±0.05 | 0.92±0.12 | 0.91±0.06 | 0.61±0.06 | **0.14±0.01** | H(5)=19.13, P=0.0018 |
| protein levels | N/A | 1.0±0.0 | 1.3±0.3 | 1.2±0.3 | 1.1±0.2 | 0.9±0.0 | H(4)=4.04, P=0.40 |
| **BC11-38** | | | | | | | |
| cAMP-PDE | **0.06±0.00** | 1.0±0.07 | 1.09±0.03 | 1.07±0.03 | 1.15±0.02 | **0.69±0.08** | F(5,18)=76.79, P<0.0001 |
| cGMP-PDE | **0.02±0.00** | 1.0±.05 | 1.07±0.10 | 1.11±0.05 | 1.13±0.02 | **0.71±0.06** | F(5,18)=59.92, P<0.0001 |
| protein levels | N/A | 1.0±0.1 | 2.4±0.9 | 1.4±0.2 | 1.6±0.4 | 2.4±0.5 | F(4,15)=1.42, P=0.2767 |
| **MLG-199** | | | | | | | |
| cAMP-PDE | **0.09±0.01** | 1.0±0.02 | not tested | 0.92±0.01 | 0.92±0.04 | **0.78±0.05** | H(4)=16.07, P=0.0029 |
| cGMP-PDE | **0.04±0.00** | 1.0±0.02 | not tested | **0.89±0.04** | **0.90±0.02** | **0.75±0.02** | F(4,15)=316.84, P<0.0001 |
| protein levels | N/A | 1.0±0.00 | not tested | 0.91±0.1 | 1.0±0.1 | 0.83±0.0 | F(3,12)=2.55, P=0.1051 |
| **MLG-185** | | | | | | | |
| cAMP-PDE | **0.09±0.00** | 1.0±0.01 | 0.94±0.03 | **0.88±0.04** | **0.33±0.04** | **0.11±0.00** | H(5)=21.28, P=0.0007 |
| cGMP-PDE | **0.01±0.00** | 1.0±0.02 | 1.02±0.08 | 0.88±0.03 | 0.56±0.21 | **0.05±0.00** | H(5)=17.93, P=0.003 |
| protein levels | N/A | 1.0±0.1 | 1.02±0.1 | 1.1±0.1 | 0.82±0.1 | 0.78±0.1 | F(4,15)=1.46, P=0.2635 |
| **MLG-122** | | | | | | | |
| cAMP-PDE | **0.07±0.01** | 1.0±0.04 | 1.01±0.03 | 1.03±0.03 | **0.73±0.03** | **0.75±0.02** | F(5,18)=201.7, P<0.0001 |
| cGMP-PDE | **0.00±0.00** | 1.0±0.06 | 0.96±0.07 | 1.0±0.05 | **0.76±0.05** | **0.79±0.01** | F(5,18)=66.73, P<0.0001 |
| protein levels | N/A | 1.0±0.1 | 0.99±0.1 | 1.2±0.2 | 0.78±0.2 | 1.5±.3 | F(4,15)=2.04, P=0.1407 |
| **SMQ-02-57** | | | | | | | |
| cAMP-PDE | **0.03±0.01** | 1.0±0.03 | 1.02±0.04 | **0.65±0.13** | **0.31±0.06** | **0.05±0.00** | H(5)=21.48, P=0.0007 |
| cGMP-PDE | **0.01±0.00** | 1.0±0.05 | **0.85±0.04** | **0.59±0.11** | **0.23±0.02** | **0.03±0.01** | H(5)=21.45, P=0.0007 |
| protein levels | N/A | 1.0±0.0 | 1.16±0.1 | 1.04±0.2 | 1.24±0.2 | 1.19±0.1 | F(4,15)=0.56, P=0.6975 |
| **SMQ-03-30** | | | | | | | |
| cAMP-PDE | **0.04±0.02** | 1.0±0.05 | 1.07±0.08 | **0.76±0.06** | **0.28±0.02** | **0.23±0.02** | F(5,18)=85.79, P<0.0001 |
| cGMP-PDE | **0.0±0.00** | 1.0±0.11 | 1.13±0.07 | 0.87±0.07 | **0.33±0.01** | **0.4±0.04** | H(5)=20.39, P=0.001 |
| protein levels | N/A | 1.0±0.1 | 1.27±0.3 | 1.89±0.4 | 2.37±0.7 | 2.47±0.8 | H(4)=3.83, P=0.4297 |
| **SMQ-03-20** | | | | | | | |
| cAMP-PDE | **0.02±0.01** | 1.0±0.01 | 1.01±0.01 | **0.94±0.01** | **0.36±0.04** | **0.13±0.01** | H(5)=21.77, P=0.0006 |
| cGMP-PDE | **0.0±0.01** | 1.0±0.02 | 0.99±0.03 | 0.93±0.02 | **0.42±0.02** | **0.18±0.02** | F(5,18)=451.33, P<0.0001 |
| protein levels | N/A | 1.0±0.2 | 1.12±0.2 | 1.08±0.3 | 0.89±0.3 | 1.23±0.7 | H(4)=0.96, P=0.9162 |

To facilitate comparing efficacy across compounds, PDE activity (CPMs-background/μg total protein) and total PDE11A4 protein expression (PDE11A4 r.o.d./Ponceau S r.o.d.) are expressed as a fold-change of WT + 0 μM (n=4 biological replicates/group/experiment). Data presented here reflect the mean ±SEM, with **bolding** of values indicating the group is significantly lower than WT + 0 μM, P<0.05-0.0001. Note that the raw values for the dose responses reported above for tadalafil ("1" in [1]), BC11-38 [2], MLG-185 ("14d" in [1]), SMQ-02-57 ("23b" in [1]), and SMQ-03-20 ("4g" in [3] were previously reported, but are included here alongside never-reported data for MLG-199 ("14k" in [1]), MLG-122 ("9a" in [1]), and SMQ-03-30 ("4h" in [3]) to facilitate relative comparisons of potency and efficacy across all compounds. GFP—emerald-green fluorescent protein, WT--GFP-PDE11A4.

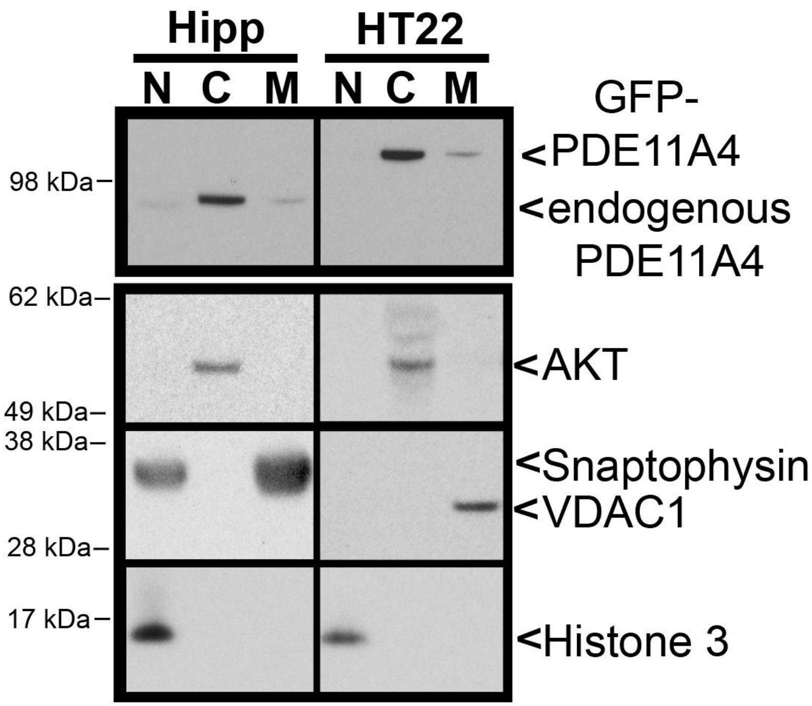


**Figure S1. Biochemical fractionation of an old mouse hippocampus and GFP-PDE11A4 transfected HT22 cells demonstrates that GFP-tagged mouse PDE11A4 expresses and subcellularly compartmentalizes in a similar manner as does endogenous mouse PDE11A4**. Biochemical fractionation was performed as previously described [4-7]. In short, cells or tissue were sonicated in ice cold fractionation buffer (20 mM Tris-HCl, pH 7.5; 2 mM MgCl2; Thermo Pierce Scientific phosphatase tablet #A32959 and protease inhibitor 3 #P0044). For those samples processed by biochemical fractionation, successive rounds of differential centrifugation were conducted. Following a low-speed spin (1000 x g) to isolate the nuclear fraction pellet, the supernatant was transferred to a new tube and subjected to a high-speed spin (89,000 x g) to isolate the cytosolic proteins in the supernatant and the membrane proteins in the pellet. The pellet was then resuspended in fractionation buffer with 0.5% Triton-X 100 to solubilize the protein, with a subsequent high-speed spin (60,000 x g) to isolate the supernatant containing soluble membrane proteins. Note that the nuclear and membrane pellets were "rinsed" prior to the resuspensions described above. That is, these pellets were resuspended in the previous buffer and respun with resulting supernatant discarded. Protein concentrations were determined by DC Protein Assay kit (Bio-Rad; Hercules, CA, USA) according to manufacturer protocol, and were subsequently equalized across samples for probing on a Western blot. Fractionation was validated by Western blots showing AKT in the cytosol, synaptophysin or VDAC1 in the membrane, and histone H3 in the nuclear fraction. The need for differing membrane markers between tissue and cells is due to the following. Synaptophysin—a marker of differentiated synaptic terminals [8]—is not detected in our HT22 cells because we are not culturing these cells using differentiating conditions. The discrepancy we see with VDAC1 is related to the greater complexity of brain tissue relative to cultured cells. In brain tissue, mitochondria are often physically tethered to larger structures and, thus, sediment during the crude, low-speed spins such as the one we use to enrich for the nuclear pellet [9-11].

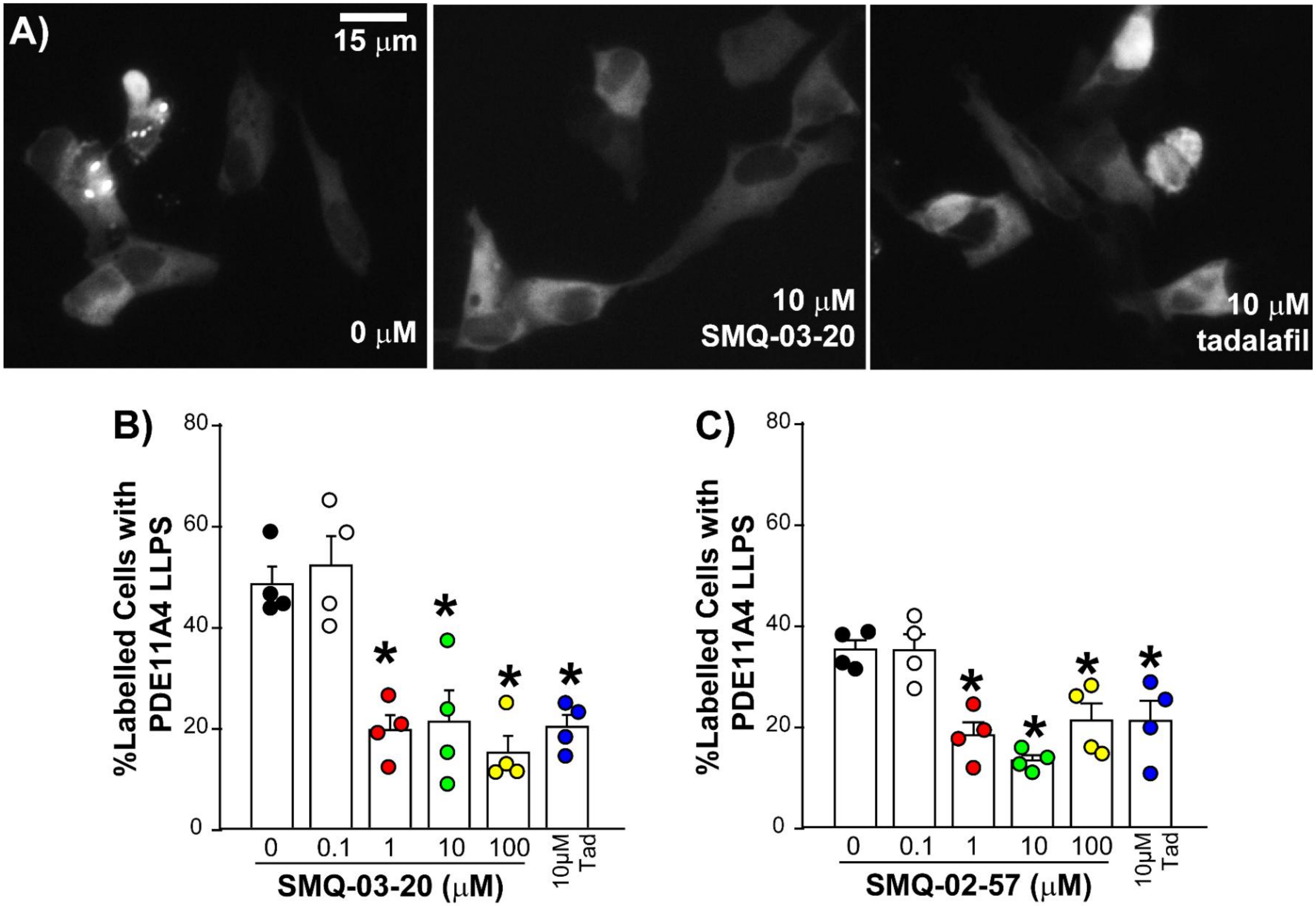


**Figure S2. PDE11Ai's reverse human PDE11A4 LLPS in human SH-SY5Y neuronal cells.** SH-SY5Y cells were transfected with hPDE11A4, allowed to form hPDE11A4 LLPS, and then treated for 1 hour with a PDE11Ai. A) Widefield microscopic images of GFP-tagged hPDE11A4 in SH-SY5Y cells treated with vehicle (i.e., 0 μM), 10 μM SMQ-03-20 or 10 μM tadalafil reveals dispersal of hPDE11A4 LLPS with PDE11Ai treatment. B) Quantification shows 1-100 μM SMQ-03-20 and 10 μM tadalafil significantly reduce hPDE11A4 LLPS ($F(5,18)=14.58$, $P<0.0001$; Post hoc vs 0 μM: 1-100μM SMQ-03-20 and 10 μM tadalafil, $P<0.001$ each). C) SMQ-02-57 also reversed hPDE11A4 LLPS in SH-SY5Y cells ($F(5,18)=10.10$, $P<0.0001$; Post hoc vs 0 μM: 1-100 μM SMQ-02-57 and 10 μM tadalafil, P=0.013-0.0006), as did MLG-185 (data not shown). Post hoc *vs 0 μM, $P<0.001$.

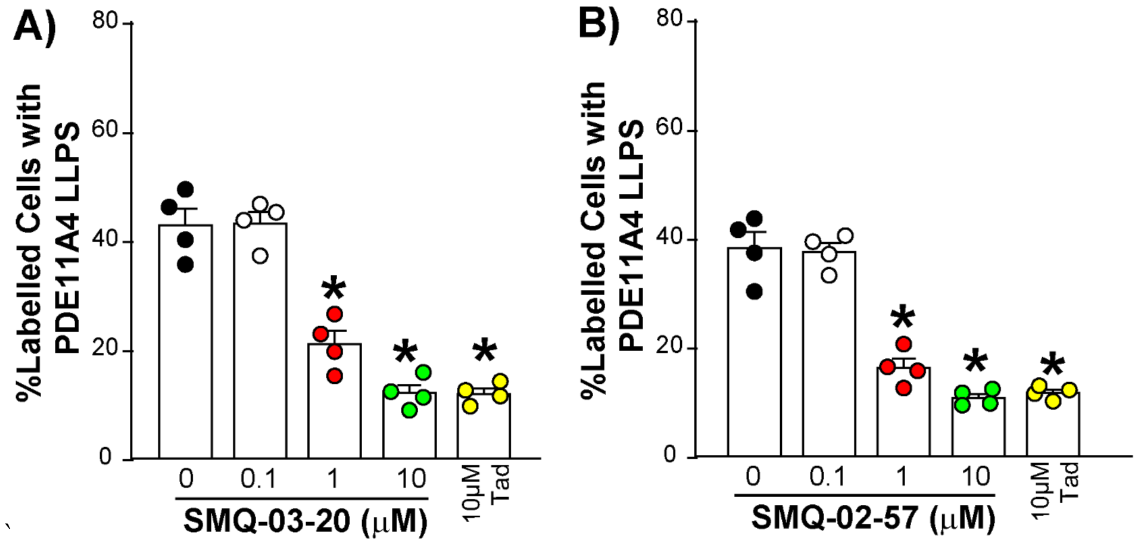


**Figure S3. PDE11Ai's prevent the formation of mouse PDE11A4 LLPS in mouse HT22 hippocampal neuronal cells.** PDE11Ai's were applied to HT22 cells following transfection with mPDE11A4, but before mPDE11A4 LLPS took place. A) 1-10 μM SMQ-0320 and 10 μM tadalafil ($F(4,15)=55.14$m $P<0.0001$; Post hoc vs 0 μM: 1-10 μM SMQ-03-20 and 10 μM tadalafil, $P=0.0002$ for each) as well as B) 1-10 μM SMQ-02-57 ($F(4,15)=63.91$, $P<0.0001$; Post hoc vs 0 μM: 1-10 μM SMQ-03-20 and 10 μM tadalafil, $P=0.0002$ for each) prevented the formation of PDE11A4 LLPS, suggesting prophylaxis. Data representative of 2 experiments. Post hoc *vs 0 μM, $P<0.0002$.

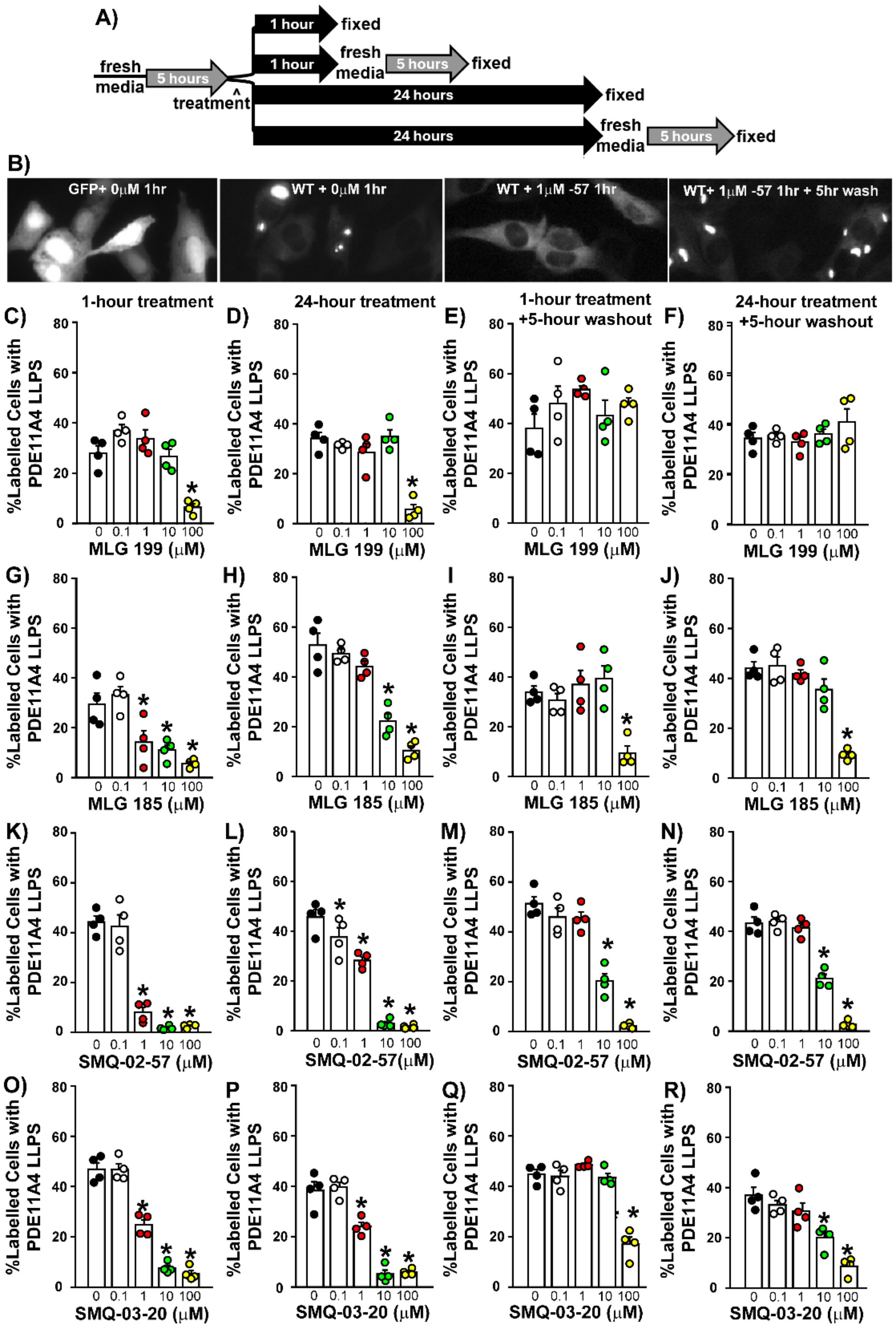


**Figure S4. 1hr and 24hr PDE11Ai treatments equally disperse PDE11A4 LLPS with some effects reversible upon washout. A)** Graphical outline of experimental time course. **B)** Exemplar images of HT22 neuronal cells expressing high levels of GFP alone (1st image; no droplets) or GFP-PDE11A4 following treatment with vehicle (2nd image; many droplets), a PDE11A inhibitor (3rd image), or a PDE11A inhibitor followed by a 5-hour washout (4th image; PDE11Ai shown is 1 μM SMQ-02-57). MLG-199 reversed PDE11A4 LLPS with similar efficacy and potency following either a **C)** 1-hour (ANOVA: F(4,15)=19.51, P<0.0001; Post hoc: 0 vs. 100 μM, P=0.0003) or **D)** 24-hour treatment (ANOVA: F(4,15)=23.85, P<0.0001; Post hoc: 0 vs. 100 μM, P=0.0002). PDE11A4 again formed droplets (i.e., de-mixed) following washout of the **E)** 1-hour (ANOVA: F(4,15)=1.34, P=0.3006) and **F)** 24-hour MLG-199 treatments (failed equal variance; ANOVA on Ranks: H(4)=1.58, P=0.8122). MLG-185 reversed PDE11A4 LLPS to a similar extent with either a **G)** 1-hour (ANOVA: F(4,15)=12.51, P=0.0001; Post hoc vs 0 μM: 1 μM P=0.0066, 10 μM P=0.0045; 100 μM P=0.001) or **H)** 24-hour treatment (fails equal variance; ANOVA on Ranks: H(4)=15.84, P=0.0032; Post hoc: 0 vs 10-100 μM, P<0.05 for each). PDE11A4 de-mixed after washing out the 1-10 μM, but not 100 μM, **I)** 1-hour treatment (ANOVA: F(4,15)=8.95, P=0.0007; Post hoc 0 vs. 10 μM P=0.6187, 0 vs. 100 μM, P=0.0012) and **J)** 24-hour MLG-185 treatments (ANOVA: F(4,15)=27.24, P<0.0001; Post hoc 0 vs. 10 μM P=0.11, 0 vs. 100 μM, P=0.0002). SMQ-02-57 reversed PDE11A4 LLPS with similar efficacy and potency with either a **K)** 1-hour (equal variance fails; ANOVA on Ranks: H(4)=16.0, P=0.0023; Post hoc: 0 vs. 1-100 μM, P<0.05 for each) or **L)** 24-hour treatment (ANOVA: F(4,15)=76.29, P<0.0001; Post hoc vs 0 μM: 0.1 μM P=0.0268, 1 μM P=0.0004, 10 μM P=0.0002, 100 μM P=0.0001). PDE11A4 fully de-mixed after washing out 1 μM, partially de-mixed after washing out 10 μM, and failed to de-mix after washing out 100 μM of the **M)** 1-hour (ANOVA: F(4,15)=61.43, P<0.0001; Post hoc vs. 0 μM: 0. 1 μM P=0.1755, 1 μM P=0.2938, 10 μM, P=0.0002, , P=0.0001) and N) **24-hour** SMQ-02-57 treatments (ANOVA F(4,15)=111.29, P=0.0001; Post hoc vs. 0 μM: 0. 1 μM P=0.8258, 1 μM P=0.463, 10 μM and 100 μM P=0.0002). Finally, SMQ-03-20 reversed

PDE11A4 LLPS with the same efficacy and potency with either a **O)** 1-hour (ANOVA: $F(4,15)=108.08$, $P<0.0001$; Post hoc: 0 vs. 1-100 μM, P=0.0002 for each) or **P)** 24-hour treatment (ANOVA: $F(4,15)=65.10$, $P<0.0001$; Post hoc vs. 0 μM: 1 μM P=0.0004, 10 and 100 μM P=0.0002 for each). **Q)** After a 1-hour treatment with SMQ-03-20, PDE11A4 fully de-mixed after washing out 1-10 μM, but not 100 μM (ANOVA: $F(4,15)=40.98$, $P<0.0001$; Post hoc vs 0 μM: 1 μM P=0.2135, 10 μM P=0.8795, 100 μM P=0.0002). **R)** Following a 24-hour SMQ-03-20 treatment, PDE11A4 demixed fully only after 1 μM, with only partial demixing after 10 μM and no demixing after 100 μM washout (ANOVA: $F(4,15)=20.19$, $P<0.0001$; Post hoc vs. 0 μM: 1 μM P=0.2159, 10 μM, P=0.0017, 100 μM, P=0.0002). *vs 0 μM <0.05-0.0001. Data plotted mean ±SEM.

**Table S2. Treatment of GFP-mPDE11A4 transfected HT22 cells with a cAMP or cGMP analogue was not sufficient to reduce PDE11A4 LLPS suggesting that the ability of PDE11A4 inhibitors to reduce PDE11A4 LLPS is not related to their ability to increase cAMP/cGMP as a result of inhibiting PDE11A4 catalytic activity.**

| | **0 μM** | **0.1 μM** | **1 μM** | **10 μM** | **100 μM** | **10 μM tadalafil (+control)** | **Effect of treatment** |
|---|---|---|---|---|---|---|---|
| **8-Br-cAMPs** mean: | 26.49 | 24.62 | 24.86 | 29.09 | 26.80 | **6.28** | $F(5,18)=25.65$, $P<0.0001$ |
| ± SEM: | 2.47 | 1.31 | 1.63 | 1.15 | 1.57 | **1.45** | *post hoc*: tadalafil vs all, P<0.0002 |
| **Sp-5,6-DCI-cBIMPs** mean: | 35.58 | 28.58 | 36.84 | 36.03 | 35.82 | **5.15** | $F(5,18)=30.6$, $P<0.0001$ |
| ± SEM: | 1.33 | 3.90 | 0.88 | 2.71 | 2.02 | **0.96** | *post hoc*: tadalafil vs all, P<0.0002 |
| **8-Br-cGMPs** mean: | 27.99 | 28.68 | 30.08 | 28.84 | 30.60 | **8.45** | $F(5,18)=22.65$, $P<0.0001$ |
| ± SEM: | 1.84 | 1.57 | 1.93 | 1.28 | 2.31 | **1.65** | *post hoc*: tadalafil vs all, P<0.0002 |
| **Sp-8pCPT-cGMPs** mean: | 33.44 | 36.38 | 33.39 | 37.57 | 35.03 | **4.99** | $F(5,18)=36.33$, $P<0.0001$ |
| ± SEM: | 2.05 | 2.68 | 1.86 | 2.81 | 1.22 | **1.13** | *post hoc*: tadalafil vs all, P<0.0002 |

Data shown reflects the %labelled cells exhibiting PDE11A4 LLPS following treatment with 0-100 μM of a cAMP or cGMP analogue or 10 μM tadalafil as a positive control. Bolded values P<0.0002 vs vehicle (0 μM).

**Table S3. Neither cAMP nor cGMP blockers prevent PDE11Ai's from dispersing PDE11A4 LLPS suggesting that this dispersal occurs independently of inhibitor-induced increases in cAMP/cGMP levels.**

| | vehicle + 0 μM | vehicle + 100 μM | BC11-38 + 0 μM | BC11-38 + 100 μM | Two Way ANOVA |
|---|---|---|---|---|---|
| **Rp-8-Br-cAMP** mean: | 60.95 | 65.95 | **30.96** | **37.07** | *effect of Rp:* F(1,12)=2.41, P=0.1466 |
| ± SEM: | 4.18 | 3.72 | **3.94** | **2.10** | *effect of PDE11Ai:* F(1,12)=67.50, P<0.0001 |
| **Rp-cAMP mean:** | 64.49 | 67.66 | **27.76** | **33.15** | *effect of Rp*: F(1,12)=1.74, P=0.2121 |
| ± SEM: | 4.78 | 3.12 | **2.00** | **2.35** | *effect of PDE11Ai*: F(1,12)=120.32, P<0.0001 |
| **Rp-8-pCPT-cGMPs** mean: | 65.18 | 65.67 | **19.78** | **28.70** | *effect of Rp:* F(1,12)=1.48, P=0.2464 |
| ± SEM: | 2.28 | 1.79 | **5.78** | **4.22** | *effect of PDE11Ai:* F(1,12)=113.73, P<0.0001 |

Data shown reflects the %labelled cells exhibiting PDE11A4 LLPS following treatment with 0 or 100 μM BC11-38 in combination with 0 or 100 μM a cAMP (Rp-8-Br-cAMP, Rp-cAMP) or cGMP blocker (Rp-8-pCPT-cGMP). Results replicate our previously published findings showing that neither cAMP nor cGMP blockers interfere with the ability of PDE11Ai's to disperse PDE11A4 LLPS. **Bolded** values P<0.0001 vs vehicle (0 μM).

**Table S4.** Phosphodiesterase activity and total PDE11A4 protein expression in ventral hippocampus of young and old vehicle-treated and SMQ-03-20-treated mice.

| | | **Young Vehicle** | **Young SMQ-03-20** | **Old Vehicle** | **Old SMQ-03-20** | **Two-Way ANOVA:** |
|---|---|---|---|---|---|---|
| **cAMP-PDE activity (CPMs/ug total protein)** | Mean ±SEM: | 1343.33 ±230.09 | 899 ±262.03 | 1714.8 ±186.1 | 1810.2 ±106.53 | *effect of treatment:* F(1,14)=0.796, P=0.3875*;* *effect of age:* F(1,14)=10.752, P=0.0055 |
| **PDE11A4 /Poncea S Stain (r.o.d.)** | Mean ±SEM: | 1.00 ±0.20 | 1.00 ±0.08 | 1.00 ±0.24 | 0.86 ±0.14 | *effect of treatment:* F(1,14)=0.154, P=0.701; effect of age: F(1,14)=0.155, P=0.6999 |

young n=4/treatment; old n=5/treatment. r.o.d.--relative optical density

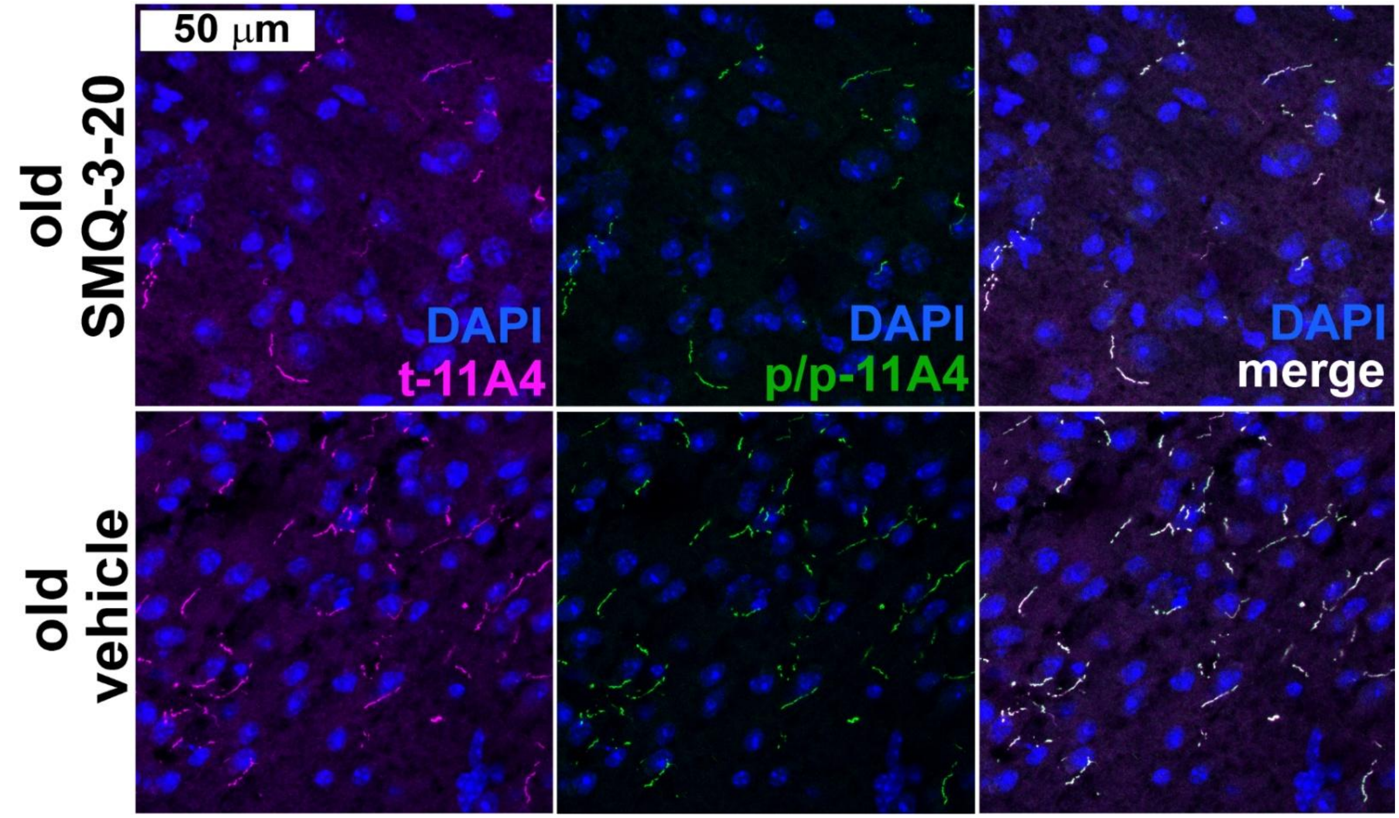


**Figure S5. An antibody recognizing pS117/pS124-PDE11A4 (p/p-11A4) labels all ghost axons that are detected using a total PDE11A4 antibody (t-11A4)** in both vehicle-treated and SMQ-3-20-treated old NIA C57BL6 mice, consistent with our previous report [7].

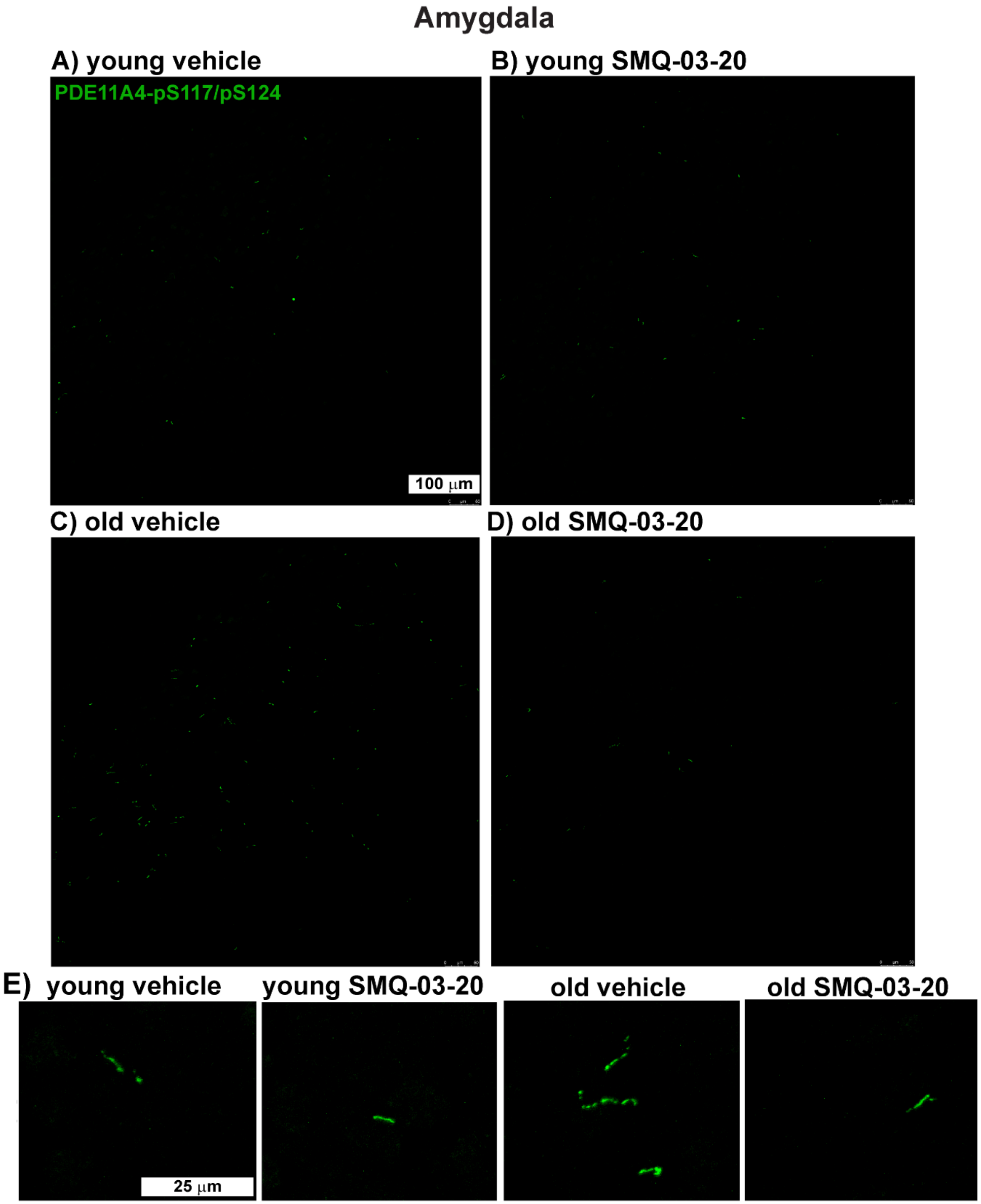


**Figure S6. Exemplar images of ghost axons in the amygdala of young versus old NIA C57BL6 mice treated with either vehicle or 30 mg/kg SMQ-03-20.** A-D) Entire image shown at low magnification. E) High magnification reveals ghost axons are frequently trails of droplets as opposed to a continuous strand.

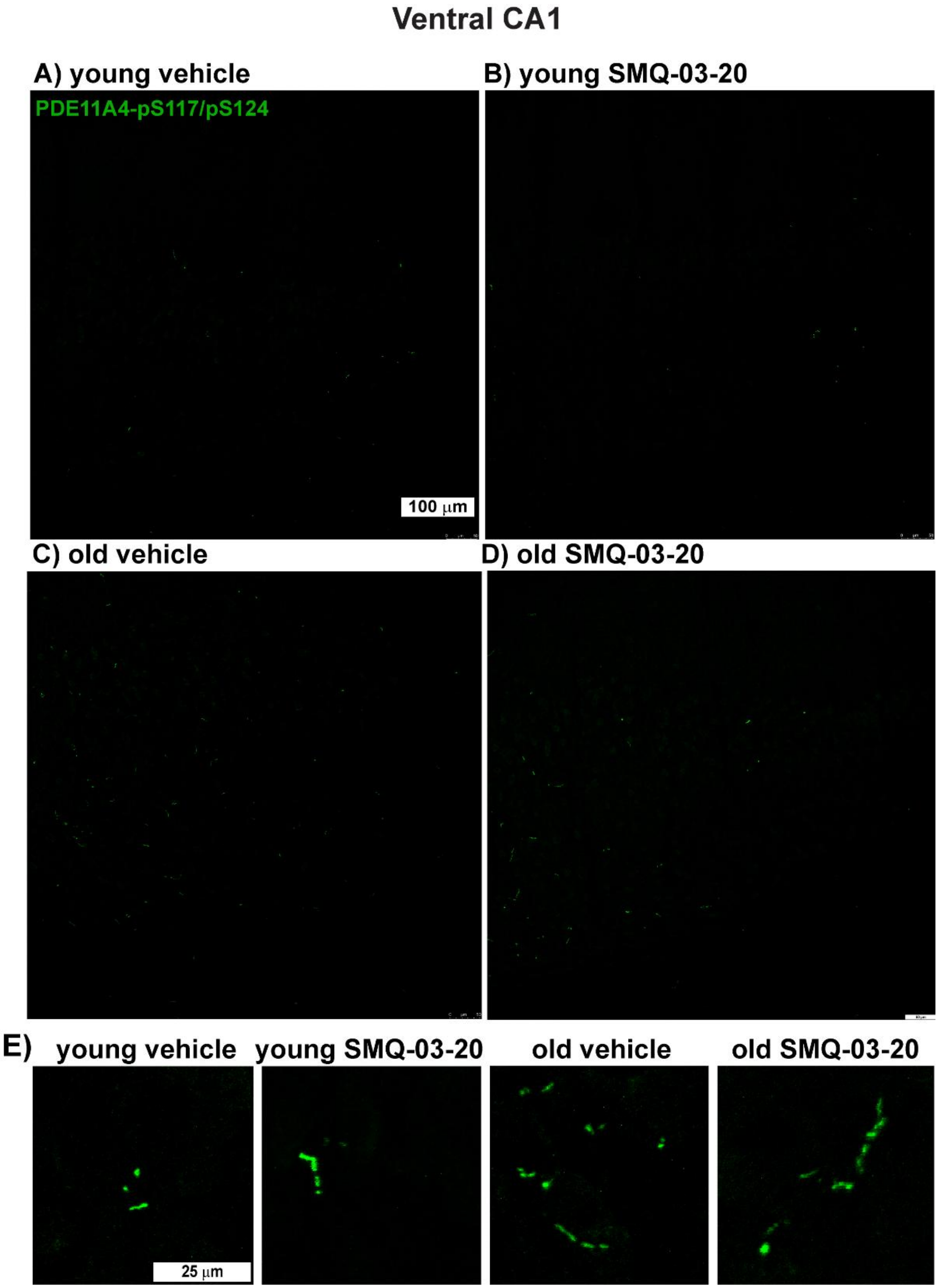


**Figure S7. Exemplar images of ghost axons in the ventral CA1 of young versus old NIA C57BL6 mice treated with either vehicle or 30 mg/kg SMQ-03-20.** A-D) Entire image shown at low magnification. E) High magnification reveals ghost axons are frequently trails of droplets as opposed to a continuous strand.

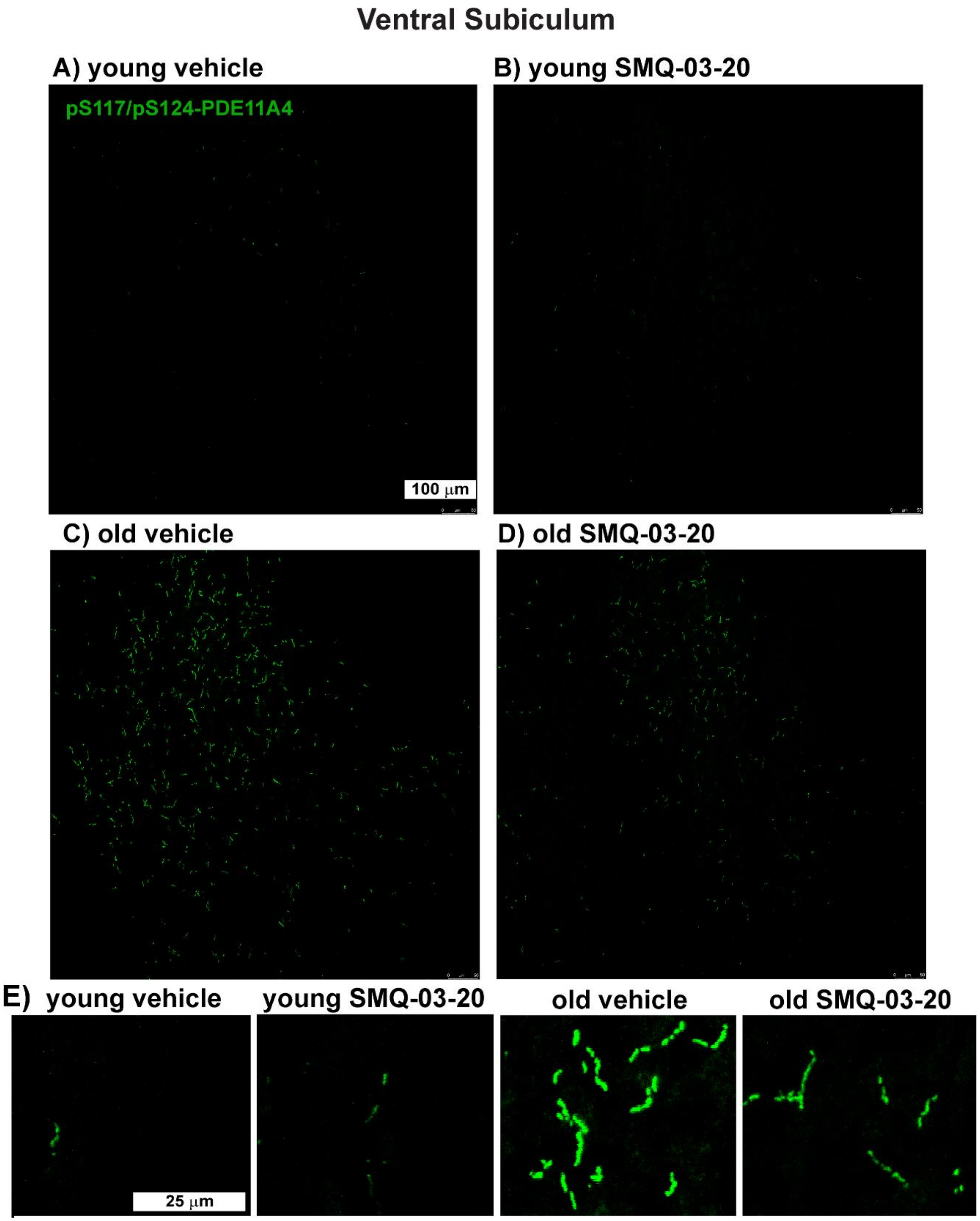


**Figure S8. Exemplar images of ghost axons in the ventral subiculum of young versus old NIA C57BL6 mice treated with either vehicle or 30 mg/kg SMQ-03-20.** A-D) Entire image shown at low magnification. E) High magnification reveals ghost axons are frequently trails of droplets as opposed to a continuous strand. The ventral subiculum is the brain region with the densest expression of PDE11A4 ghost axons, making them clearly visible even at low magnification

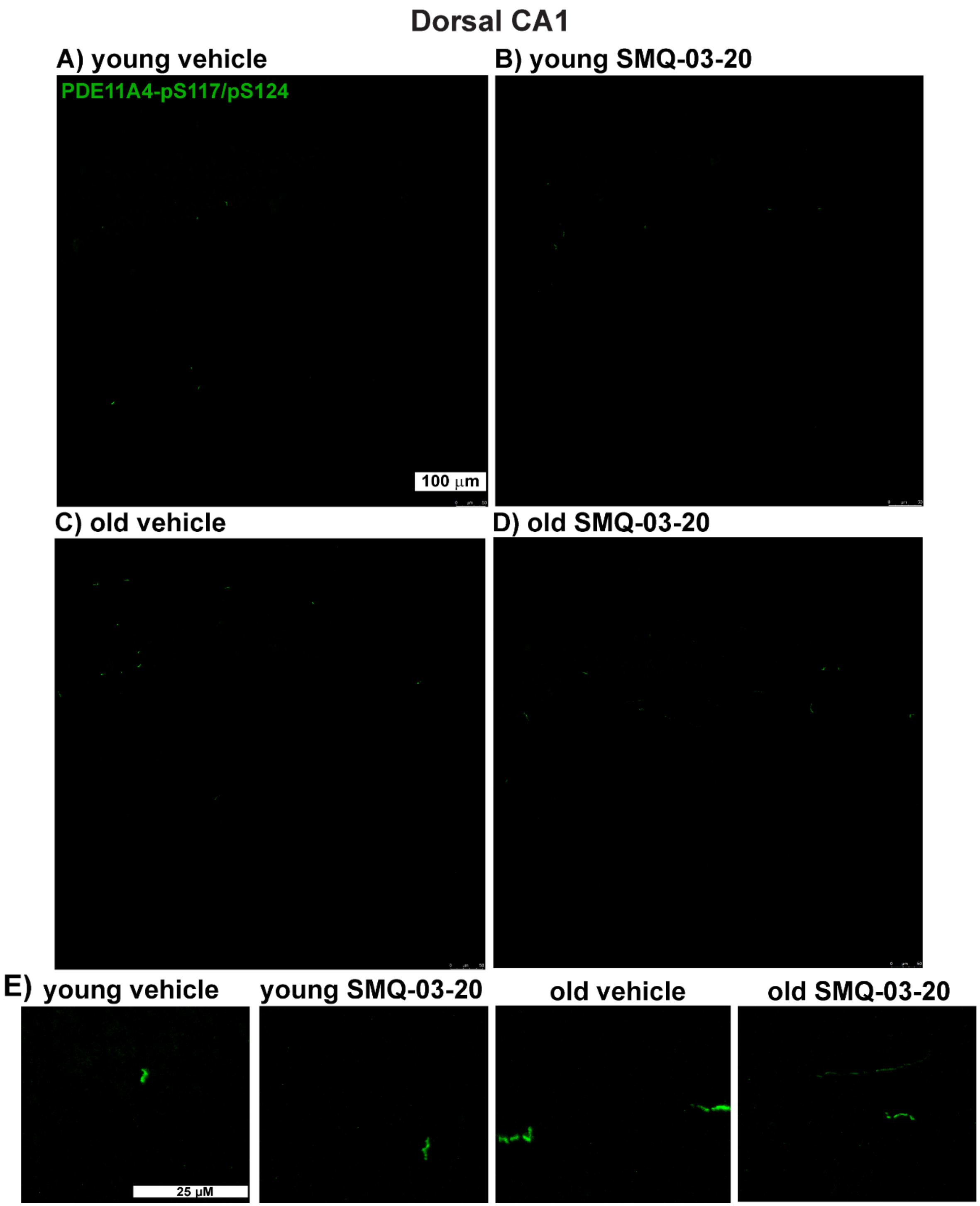


**Figure S9. Exemplar images of ghost axons in the dorsal CA1 of young versus old NIA C57BL6 mice treated with either vehicle or 30 mg/kg SMQ-03-20.** A-D) Entire image shown at low magnification. E) High magnification reveals ghost axons are frequently trails of droplets as opposed to a continuous strand.

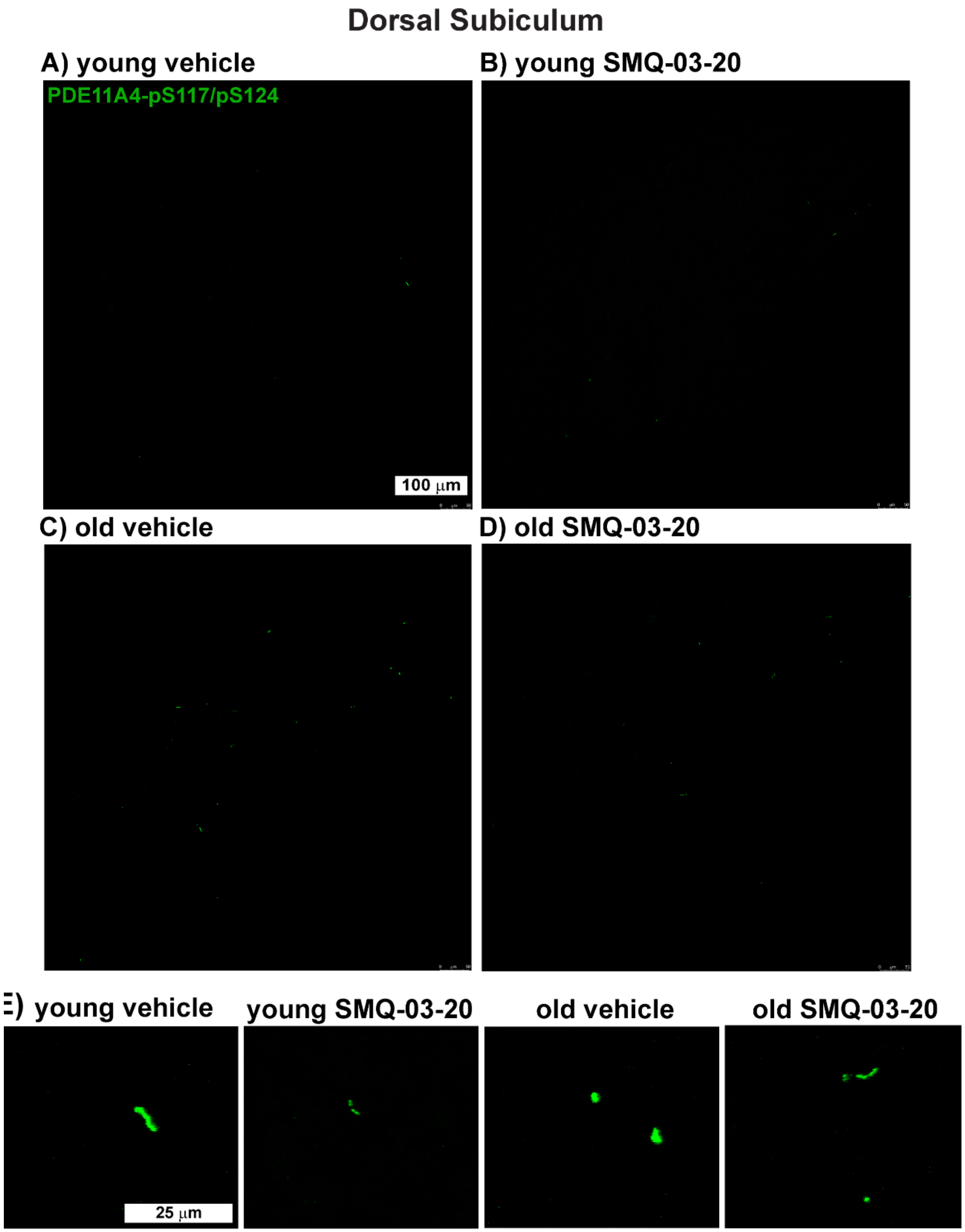


**Figure S10. Exemplar images of ghost axons in the dorsal subiculum of young versus old NIA C57BL6 mice treated with either vehicle or 30 mg/kg SMQ-03-20.** A-D) Entire image shown at low magnification. E) High magnification images.

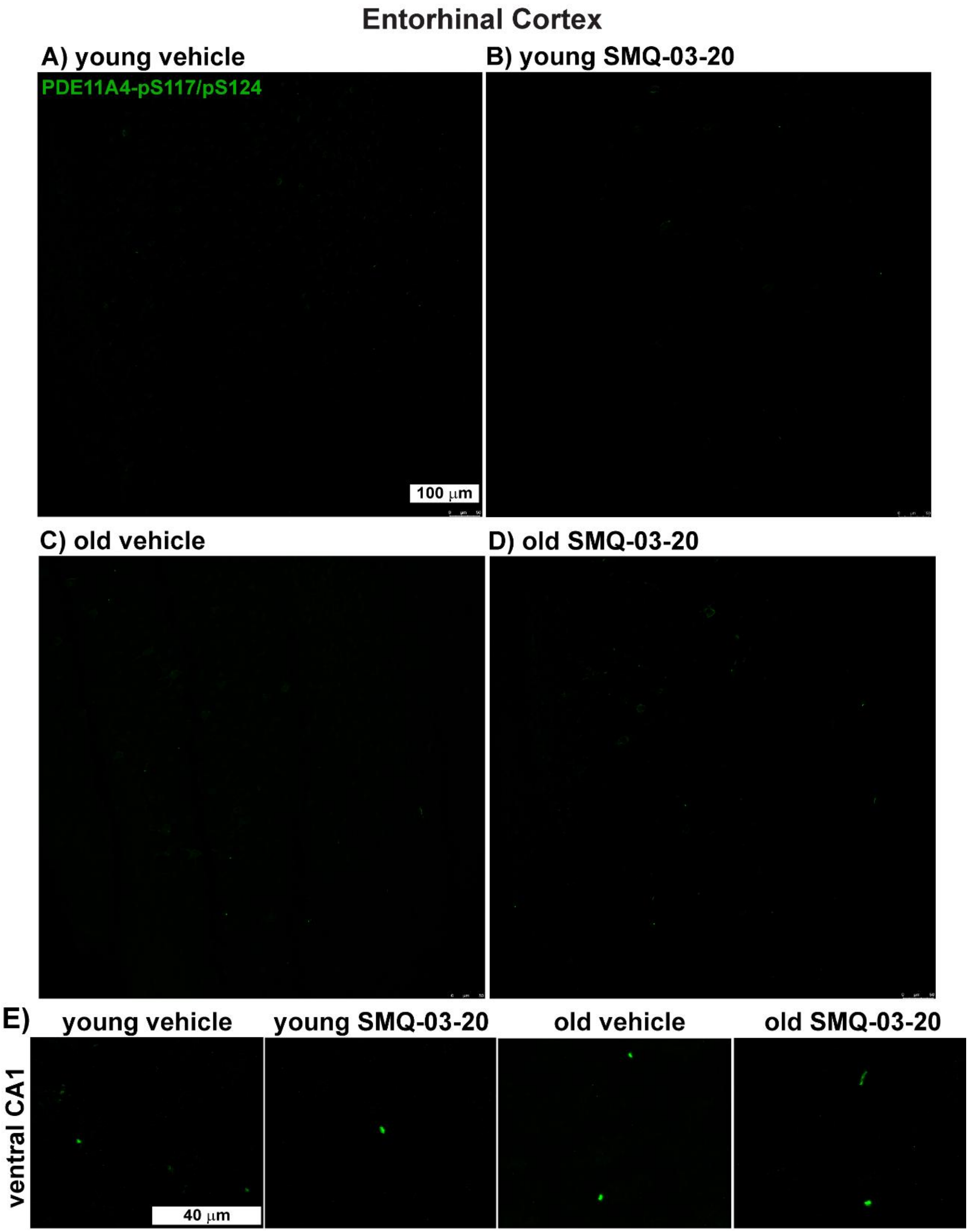


**Figure S11. Exemplar images of ghost axons in the entorhinal cortex of young versus old NIA C57BL6 mice treated with either vehicle or 30 mg/kg SMQ-03-20.** A-D) Entire image shown at low magnification. E) High magnification images. Note that ghost axons in the cortex appear shorter and stubbier than those found in the hippocampus and amygdala perhaps reflecting a differential orientation of these structures (i.e., oriented more so to the caudal or horizontal plane as opposed to the sagittal plane of sectioning).

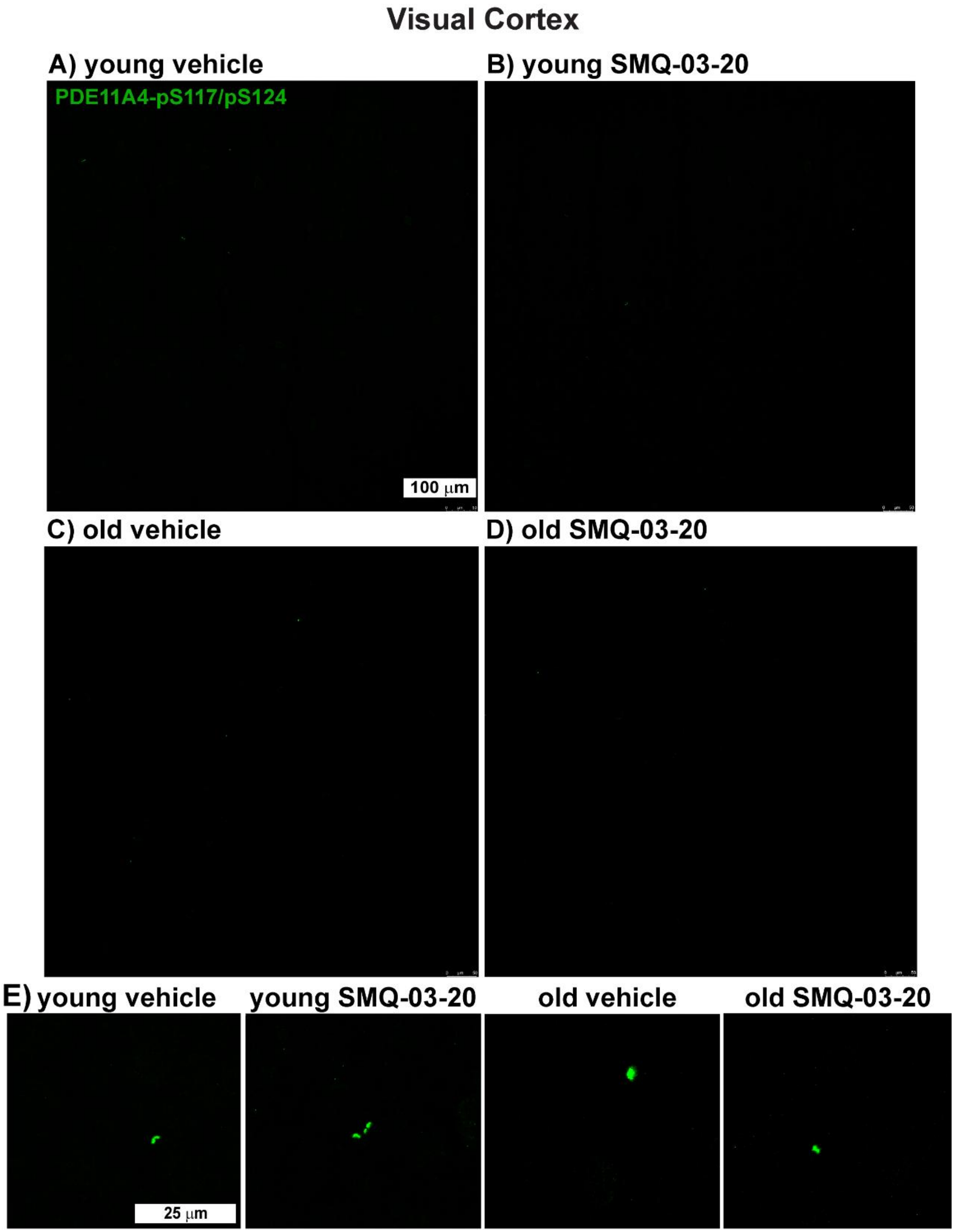


**Figure S12. Exemplar images of ghost axons in the visual cortex of young versus old NIA C57BL6 mice treated with either vehicle or 30 mg/kg SMQ-03-20.** A-D) Entire image shown at low magnification. E) High magnification images. Ghost axons are rarely found in this region and appear to be the shortest of all regions perhaps reflecting a differential orientation of these structures (i.e., oriented to the caudal plane as opposed to the sagittal plane of sectioning).

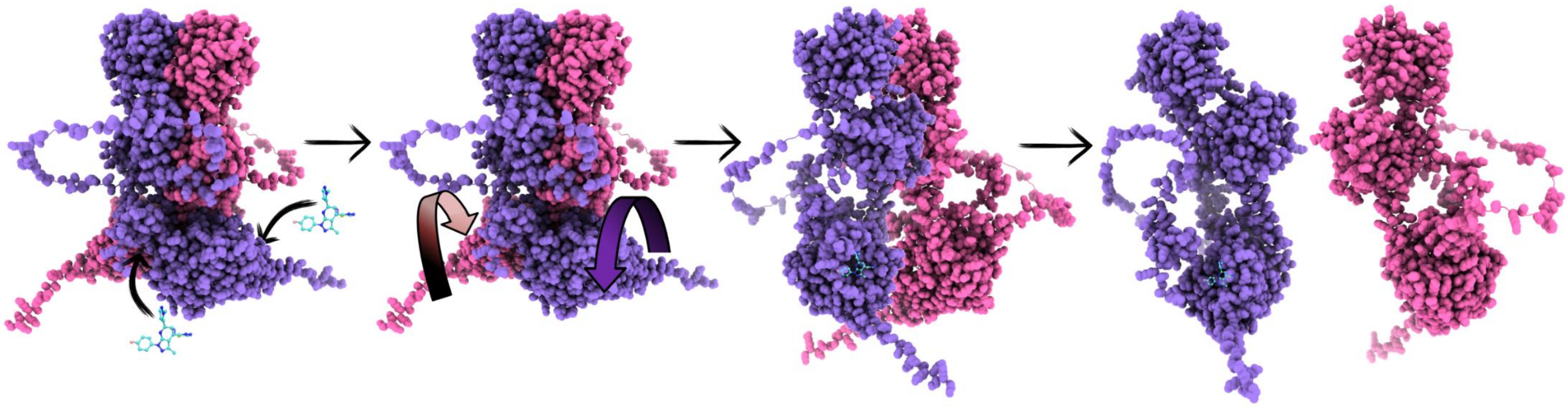

**Figure S13. Schematic of working hypothesis.** PDE11Ai's reduce LLPS, at least in part, because binding of PDE11Ai's to the catalytic pocket triggers a conformational change (e.g., a rotation) that ripples through the protein structure to weaken multivalent interactions at the GAF-B binding domain and ultimately reduce homodimerization of the protein. Our hypothesis is consistent with the fact that binding of select PDE4 inhibitors to the catalytic domain of PDE4A4 induces a conformational change that transmits to the N-terminal portion of the enzyme to alter subcellular localization (i.e., "inside-out" signaling) [12]. AlphaFold3 [13] used to predict conformation of PDE11A4 homodimer and monomer and ChimeraX 1.11.1 [14] was used to generate the graphical rendering.

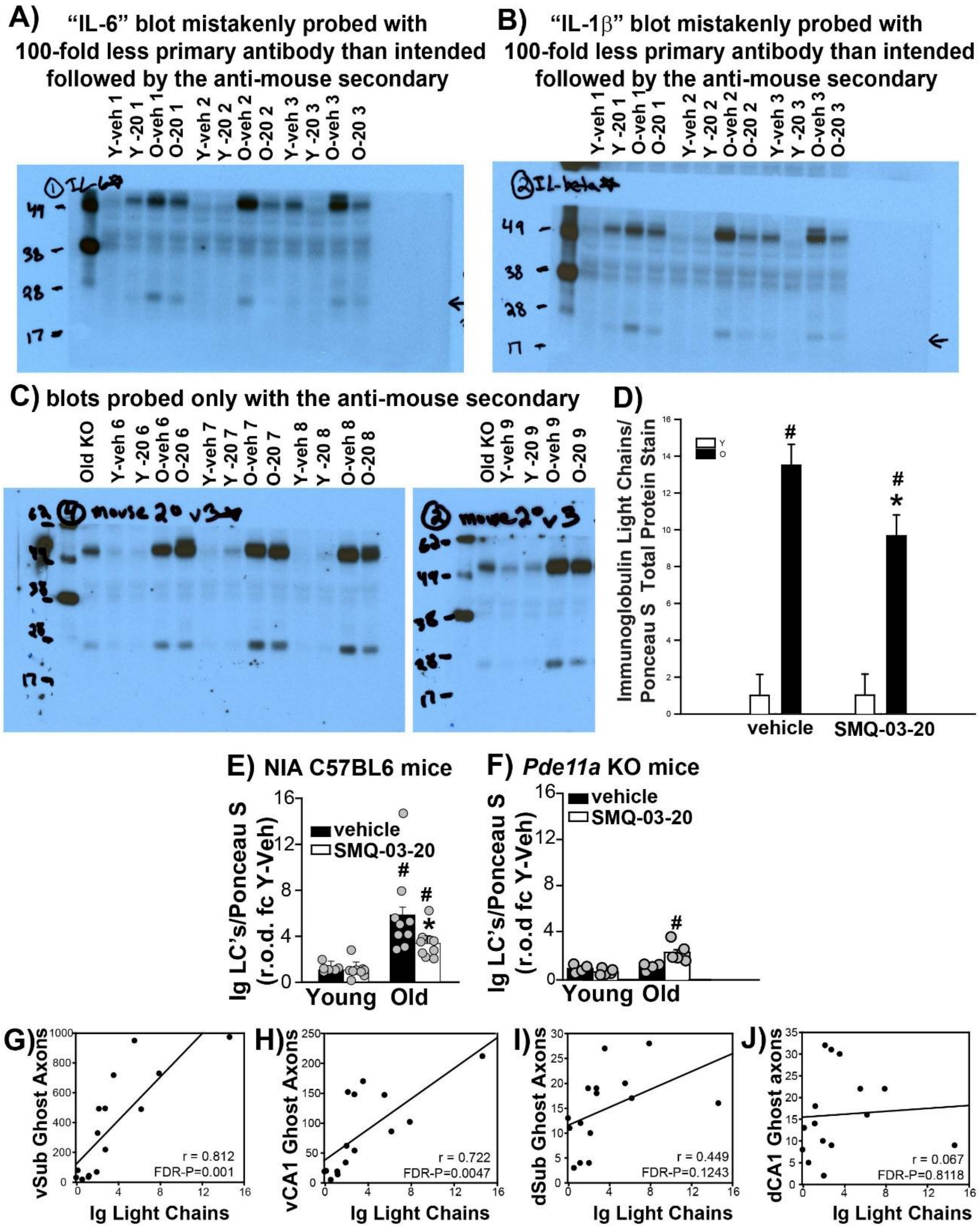


**Figure S14. Re-evaluation of blots originally intended to measure cytokines identifies age-related accumulation of immunoglobulin light chains.** A) Representative blot from the experiment originally intended to measure IL-6. Due to a dilution miscalculation, the IL-6 primary antibody was applied at a 1:80,000 dilution, corresponding to a 100-fold lower antibody concentration than intended. B) Representative blot of the same sample set from the experiment originally intended to measure IL-1β, in which the primary antibody was similarly applied at a 1:300,000 dilution. The highly similar banding patterns in A and B suggested that the detected signal arose predominantly from the anti-mouse secondary antibody rather than the intended cytokine targets. C) To test this interpretation, the remaining NIA C57BL6 samples were probed with anti-mouse secondary antibody alone. The secondary-only blots reproduced the characteristic banding pattern observed on the original "IL-6" and "IL-1β" blots, including the band originally attributed to the cytokines (black arrows), demonstrating that this signal was generated by secondary-antibody detection of endogenous mouse immunoglobulin. Based on its molecular weight, this band corresponds to immunoglobulin light chains (Ig LCs). D) Densitometric quantification of the secondary-only control blots (n=4/group) showed the same qualitative pattern across groups as observed in the original full-cohort "IL-6" and "IL-1β" blots. However, this small sample set was underpowered (n=4/group; Power with $\alpha = 0.05$ for age x treatment : 0.228483). As such, formal statistical analyses were applied only to the original full-cohort "IL-6" blots (main text Figure 8) and "IL-1β" blots. E) Quantification of Ig LCs from the original full-cohort "IL-1β" blots from

NIA C57BL6 mice showed effects consistent with those obtained from the original “IL-6” blots reported in Figure 8 (failed equal variance; RST for effect of age within vehicle: T(7,9)=28.0, FDR-*P*=0.004; effect of age within SMQ-03-20: T(9,9)=49.0, FDR-*P*=0.003; effect of SMQ-03-20 within old: T(9,9)=111.0, FDR-*P*=0.0364; effect of SMQ-03-20 within young: T(7,9)=69.0, FDR-*P*=0.3408). F) Quantification of Ig LCs from the original “IL-1β” blots of samples from *Pde11a* KO mice (2-way ANOVA, age × treatment: F(1,14)=10.25, *P*=0.0064; post hoc effect of age: vehicle-treated, *P*=0.6817; SMQ-03-20-treated, *P*=0.0003). G–J) Correlations between Ig LC levels quantified from the original “IL-1β” blots and PDE11A4 ghost axon levels in G) ventral subiculum (vSub; r=0.812, FDR-*P*=0.001), H) ventral CA1 (vCA1; r=0.722, FDR-*P*=0.0063), I) dorsal subiculum (dSub; r=0.449, FDR-*P*=0.1243), and J) dorsal CA1 (dCA1; r=0.067, FDR-*P*=0.8118). *P* values indicated by symbols: * vs old vehicle, *P*≤0.0364; # vs young, *P*≤0.004.
